# Multiple Myeloma associated DIS3 mutations drive AID-dependent IGH Translocations

**DOI:** 10.1101/2023.07.27.550610

**Authors:** Tomasz M. Kuliński, Olga Gewartowska, Mélanie Mahé, Karolina Kasztelan, Janina Durys, Anna Stroynowska-Czerwińska, Marta Jedynak-Slyvka, Ewelina P. Owczarek, Debadeep Chaudhury, Marcin Nowotny, Aleksandra Pękowska, Bertrand Séraphin, Andrzej Dziembowski

## Abstract

Role of dominant DIS3 mutations in multiple myeloma (MM) remains elusive. These mutations decrease the exoribonucleolytic activity of DIS3, a key nuclear RNA-degrading enzyme. Utilizing knock-in mice with clinical Dis3 G766R variant, we demonstrate an increased frequency of aberrant chromosomal translocations in B-cells, leading to plasmacytoma, an early-stage MM model. DIS3-dependent translocations display characteristics of aberrant activation-induced deaminase (AID) activity. In clinical MM samples with DIS3 mutations, driver genes also show AID-dependent lesions. Mechanistically, mutated DIS3 accumulates on chromatin-associated RNA substrates, including aberrant AID action sites, fostering oncogenic chromosomal rearrangements. Translocations occur during immunoglobulin class switch recombination, a process otherwise unaffected in MM patients or mice with mutated DIS3. Dis3 G766R mutation does not alter chromatin architecture in activated B-cells but hijacks it to bring together enhancers with proto-oncogenes permanently. In conclusion, gain-of-function DIS3 mutations increase nuclear exosome and AID affinity to chromatin, facilitating IGH translocations and driving MM transformation.

## INTRODUCTION

Multiple myeloma (MM) is the second most prevalent hematological malignancy among adults. It originates during the terminal differentiation of B lymphocytes into plasma cells, which reside in the bone marrow and produce massive amounts of antibodies^1,2^. MM cells, like plasma cells, have already undergone all genome editing steps associated with the maturation of activated B lymphocytes. These are activation-induced deaminase (AID) dependent immunoglobulin class switch recombination (CSR) and rounds of somatic hypermutations (SHM) diversifying antibodies^3,4^.

MM is classified based on its genetic lesions into hyperdiploid (HRD), distinguished by trisomies of several chromosomes, and non-hyperdiploid (non-HRD), characterized by chromosomal translocations^1,5^. Many translocations in MM involve the immunoglobulin heavy chain (IGH) locus on chromosome 14, leading to the overexpression of oncogenes such as *FGFR3, MMSET (NSD2), CCND3, CCND1, MYC,* and *MAF*^1,5–8^. A murine pristane-induced peritoneal plasmacytoma (PCT) models the occurrence of translocations driving neoplastic transformation as observed in MM^9^. In the BALB/c strain of mice, these chromosomal alterations typically originate at the IGH locus, directly or indirectly leading to the upregulation of the *Myc* locus^10–12^. These events are strictly dependent on the aberrant activity of AID^1^^3,14^. Furthermore, clinical data clearly indicate that AID-induced point mutations in a subset of MM driver genes are crucial for the development of human MM^15^.

The *DIS3* gene, essential for the function of every cell, is frequently mutated in MM^7,8,16^. *DIS3* encodes a catalytic subunit of the major eukaryotic nuclear ribonuclease exosome involved in the processing of stable RNA species, such as ribosomal RNA (rRNA) and degradation of cryptic transcription products, such as promoter upstream transcripts (PROMPTs) and enhancer RNA (eRNA)^17–19^. Most prominently, DIS3 protein degrades products of spurious RNA Pol II activity—low-level, unannotated, cryptic transcription that can account for up to 70% of the genome^19^. This function links the nuclear exosome to the regulation of super enhancer activity, in part by affecting chromatin topology^20^.

Mutations in *DIS3* gene disrupt posttranscriptional regulation of gene expression, resulting in reduced cell proliferation and an increase in cell death^18,21–24^. MM-associated *DIS3* mutations, which inhibit or, in the case of recurrent mutations (D479, D488, and R780), completely abolish the exoribonucleolytic activity of this enzyme, have dominant-negative effects^25^. These mutations arise early in MM progression, presumably driving MM. However, they are counter-selected in later stages of the disease due to their inhibitory effects on cell proliferation^25^.

DIS3 mutations are specific to non-HRD MM and as for now DIS3 is the only RNA exosome subunit associated with cancer^7,26,27^. This is despite the mutagenic potential of exosome depletion in various models and DNA damage often associated with dysfunctions in nuclear RNA 3’ end processing^17,28–36^. In this respect, it was recently shown that oncogenic mutations in one of the cyclin-dependent protein kinases (CDK13) lead to impairment of the function of cofactors of the nuclear RNA exosome, specifically ZC3H18 and MTR4, resulting in deficiencies in nuclear RNA surveillance and RNA exosome activity^37^.

The specific role of the exosome, particularly DIS3, in B-cells is related to CSR and SHM^38,39^, very complex processes involving a large number of transacting factors. AID deaminates cytosine residues in DNA, leading to the production of uracil and generating G:U mismatches that recruit DNA repair machinery. Uracil bases are then excised by the repair enzyme uracil-DNA glycosylase, resulting in abasic sites. This is followed by the cleavage of the DNA backbone by apurinic endonuclease, which creates single-strand breaks, that can be filled in by error-prone DNA polymerases, resulting in point mutations or indels (in SHM). Mismatches on both DNA strands lead to double-stranded DNA breaks, followed by non-homologous end joining (NHEJ) and large deletions (during CSR)^40^. Targeting AID to DNA, necessary for SHM and CSR^41,42^, involves transcription and rapid RNA degradation^43^. Target regions are generally GC-rich, contain tandemly repeated DGYW DNA sequence motifs, have the potential to form G-quadruplexes, and are prone to RNA polymerase stalling and to RNA-DNA hybrids (R-loops) formation^3,44–49^. *In vitro*, AID predominantly targets the non-template strand during transcription. *In vivo*, AID mutates both DNA strands, necessitating a special mechanism involving the exosome to displace the nascent RNA^39,50^. The full mechanistic details are not yet known, and the effect of MM-associated DIS3 point mutations in SHM and CSR has not been experimentally explored. Nevertheless, tight regulation of AID is critical, as its off-target activity is well-known to cause genetic lesions leading to B-cell lineage cancers^13,14,51,52^. Interestingly, super-enhancer regions in B-cells, which possess most of the features required by AID, are critical sites for AID recruitment and serve as hotspots for AID-mediated genomic instability^53^.

The *Igh* locus forms a topologically associating domain (TAD), which tightly regulates both the transcriptional activity of regions undergoing switch recombination and promotes deletional CSR in *cis*. In naïve B-cells, loop extrusion juxtaposes the 3’ *Igh* regulatory region (3’ RR) enhancers with the Sμ region. B-cell activation, transcription and the loading of the cohesin complex result in the generation of dynamic subdomains that directionally align the donor Sμ region with one of the downstream S regions for deletional CSR^54,55^. Upon *DIS3* KO, the chromosome architecture is abrogated, and CSR is lost^38^, adding another layer of the complexity of the DIS3 function in B-cells.

Analyses of clinical samples and MM cell lines did not reveal the exact oncogenic mechanism of *DIS3* gene mutation. Conflicting conclusions were drawn on whether this mechanism arises as a consequence of transcriptome deregulation^27^ or not^25^. Studies in cellular models, which suggested a generalized genome destabilization effect, primarily used DIS3 knockdown (KD) models and cell lines outside the B-cell lineage, failing to replicate the fact that only point mutations, rather than complete knockdowns, have been observed in MM patients^56^. Our attempts to generate mouse lines with recurrent MM *DIS3* alleles failed because of strong dominant negative effects leading to early embryonic lethality^25^. Therefore, to study the mechanistic implications of DIS3 dysfunctions in the development of B cell tumors, we decided to focus on a milder, yet clinically relevant DIS3 variant with glycine 766 mutated to arginine (*DIS3*^G766R^). *DIS3*^G766R^ is predicted to be highly pathogenic (Figure S1A), and previous experimental validation revealed that on a protein level, it leads to DIS3 protein stalling on structured RNA substrates^18,57^. Still, single-stranded RNA can be degraded^18,57^.

Herein, we have developed a DIS3^G766R^ knock-in mouse model to further our understanding of DIS3-mediated carcinogenesis in MM. Our analysis, supported by genomic studies of clinical samples, allowed us to propose a model whereby the DIS3 mutations lead to aberrant accumulation of DIS3 at sites of AID activity. The presence of DIS3 at the AID sites drives oncogenic chromosomal translocations. At the same time, and in contrast to *DIS3* loss of function mutations, MM *DIS3* variants do not abrogate CSR or affect the genome architecture.

## RESULTS

### Heterozygous MM-associated *Dis3^G^*^766^*R/+* mutation leads to subtle deregulation of non-coding regions of the genome

To study the role of *DIS3*^G766R^ in a native genomic context, we introduced the G766R in the mouse *Dis3* locus in the C57BL/B background (Figure S1B). Similar to the complete *Dis3* knockout, the homozygous Dis3^G766R^ mutation is embryonically lethal (Figure S1C) in agreement with the pathogenic character of this clinically observed DIS3 mutation. At the same time, heterozygous *Dis3*^G766R^ animals did not exhibit developmental phenotypes nor hematologic abnormalities (Figure S1D,E).

As a next step, we profiled the transcriptomes and proteomes of *in vitro* activated WT and *Dis3^G^*^766^*^R/+^* B-cells isolated from mice. Consistent with the lack of developmental phenotypes, the effect of *Dis3^G^*^766^*^R/+^* on mRNA levels was very subtle, contrasting the massive gene expression change and activation of genes related to DNA damage response upon DIS3 inactivation^18,25,56,57^. We found 16 differentially expressed genes (DEGs) in naïve, 3 DEGs in B cells upon 3 days of activation, and no DEG B-cells activated for 5 days (Figure 1A, Figure S2A, Table S1-S3). Furthermore, not a single transcript was upregulated over 2-fold. We also analyzed transcriptional responses accompanying differentiation after *in vitro* activation of B-cells and observed no significant effect of heterozygous *Dis3^G^*^766^*^R^* mutation (Figure 1B, Figure S2B, Table S4-S7). Accordingly, there were no significant changes at the proteome level of Dis3^G766R/+^ and Dis3^+/+^ primary B-cells activated for 3 days *in vitro* (Figure 1C, Table S8).

**Figure 1.**
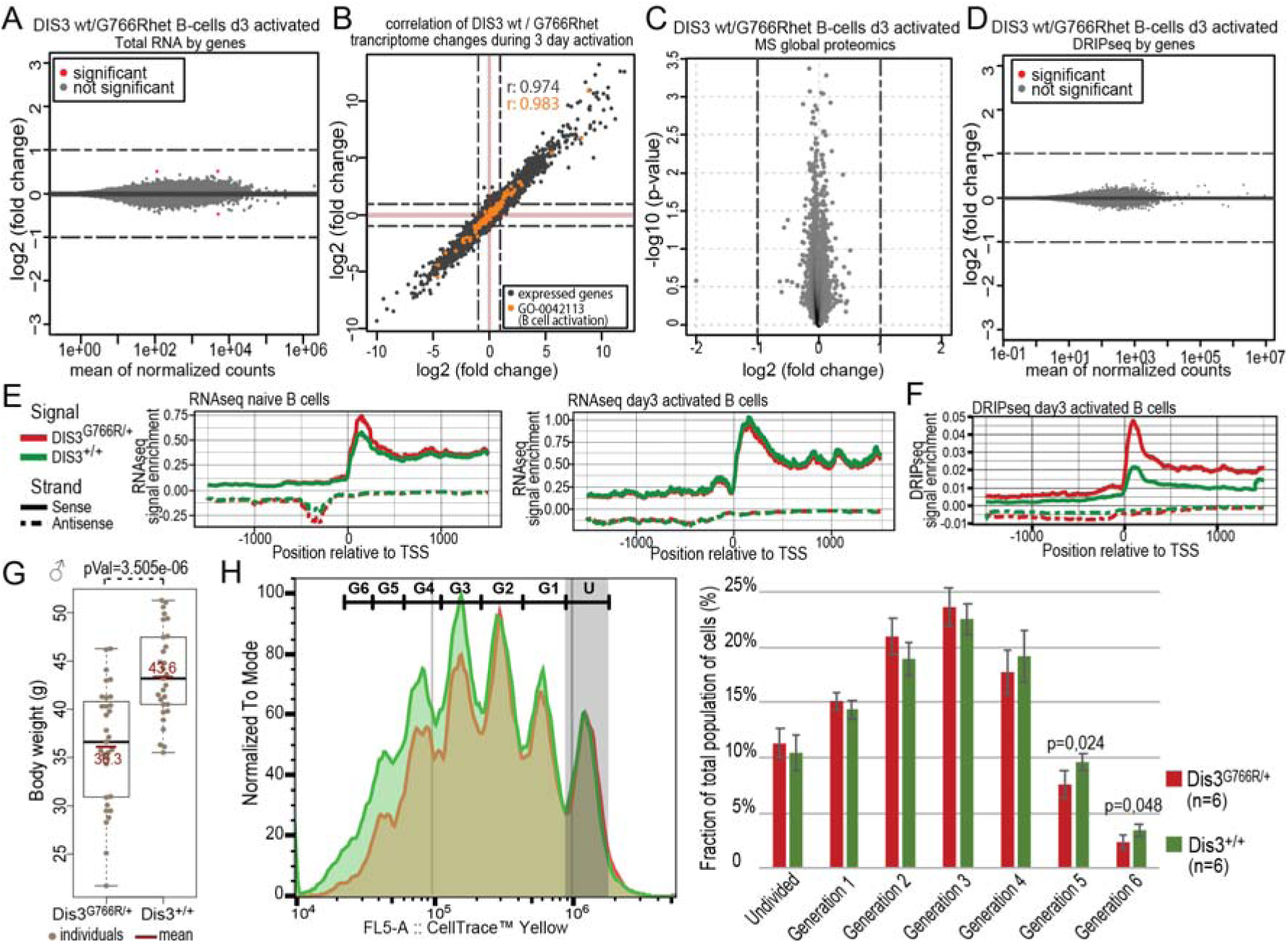
Dis3^G766R/+^ knock-in mice exhibit mild deregulation of non-coding regions. (A) MA plot comparing the transcriptome of primary Dis3^G766R/+^ and Dis3^+/+^ B-cells, *in vitro* activated for 3 days (n=3 for each genotype). (B) Correlation of transcriptome changes in the DIS3^G766R/+^ and DIS3^+/+^ during *in vitro* activation for 3 days. (C) Global proteome MS analysis of Dis3^G766R/+^ and Dis3^+/+^ primary B-cells, *in vitro* activated for 3 days (n=3 for each genotype). (D) MA plot comparing RNA/DNA hybrid DRIP-seq signal (with subtracted RNase H control signal) over annotated transcriptome Dis3^G766R/+^ and Dis3^+/+^ B-cells, *in vitro* activated for 3 days (n=2 for each genotype). Dashed lines on all plots represent a threshold of 2-fold change. (E) Meta-analyses of RNA-seq signal over TSS of expressed genes in naïve and day 3 activated B-cells. (F) Meta-analyses of DRIP-seq signal over TSS of expressed genes in day 3 activated B-cells. (G) Body weight male WT and heterozygous littermate Dis3^G766R/+^ mice aged 30- to 50-weeks. (H) Representative examples of CellTrace™ Yellow proliferation assay performed on splenic B-cells activated *in vitro* for 3 days. Quantification of CellTrace™ Yellow proliferation assay performed on B-cells activated *in vitro* for 3 days, with 6 biological replicates per genotype. Statistical analysis was conducted using a two-tailed t-test.

Given the limited effect of Dis3^G766R/+^ on the steady-state transcriptome we aimed to test whether RNA/DNA hybrids accumulate in exosome-deficient cells, potentially destabilizing the genome. For this reason, we used DNA/RNA hybrids immunoprecipitation coupled with high-throughput RNA-sequencing (DRIP-seq). Comparing the DRIP-seq over the exons of the annotated transcriptome of *Dis3^G^*^766^*^R/+^* and *Dis3^+/+^*B-cells, we scored virtually no effect of the mutation (Figure 1D).

Major DIS3 substrates represent non-protein coding RNA species with massive (up to 100-fold) accumulation of PROMPTs at the steady state of DIS3 dysfunctional cells^19^. In the case of Dis3^G766R/+^ B-cells compared to WT, no accumulation was detected when looking at individual ncRNA species. However, global analysis of the RNA-seq signal centered over active transcriptional start sites (TSS) revealed focal enrichments in regions corresponding to RNA Pol II promoter-proximal pausing and upstream regions from which PROMPTs originate (Figure 1E and S2D,E). Notably, in *Dis3^G766R/+^* B-cells, the accumulation of R loops in the promoter-proximal pausing region and over PROMPTs is far more prominently visible in the DRIP-seq (Figure 1F), indicating a very transient but detectable effect on transcriptome consistent with DIS3 role. Similar transcriptomic effects were also observed for embryonic fibroblasts (MEFs) (Figure S2F,G Table S8).

Interestingly, subtle effects of the Dis3^G766R/+^ on transcriptome have some impact on the physiology of mice. In contrast to the heterozygous *Dis3* knockout mice, which do not differ in size from WT, *Dis3*^G766R/+^ animals were smaller than their WT littermates (Figure 1G, Figure S3A-D). This reduced size corresponds to a negative impact on cell proliferation both in activated B-cells and MEFs (Figure 1H, Figure S3E,F), aligning with a conserved phenotype of DIS3 dysfunction, which includes reduced mitotic progression and increased cell death.^18,21–24^

We conclude that primary B cells do not show any detectable defects on the annotated protein-coding transcriptome or proteome. Instead, the molecular phenotype is limited to unstable R-loop forming on non-protein coding RNA species vulnerable to the biochemically mild heterozygous DIS3^G766R^ variant.

### MM DIS3 variants do not affect class switch recombination and genome architecture in B-cells

The nuclear exosome has been demonstrated to be involved in CSR. Both the dysfunction of the nuclear exosome complex^39^ and *Dis3* KO^38^ abrogate CSR. Therefore, we wanted to analyze the effect of *DIS3^G766R/+^* mutation on CSR using *in vitro*-activated primary mouse B-cells. As revealed by cytometric analysis, on average 20% of cells have undergone the switch to the IgG1 heavy chain class at day 3 post-activation in both genotypes, proving that heterozygous DIS3^G766R^ mutation does not affect CSR (Figure 2A, Figure S3G). Furthermore, immunization of mice with two anti-COVID19 vaccines (mRNA and protein-based) shows unaltered production of specific α-Spike IgG1 antibodies, proving unaffected IgM to the isotype IgG CSR as well as SHM (Figure 2B). This demonstrates the proper functioning of humoral response in *Dis3^G766R/+^*mice *in vivo*. Then we asked about the CSR status of MM samples with DIS3 mutations taking advantage of the CoMMpas MM genomic datasets. Like in the mouse model, CSR is unaffected in clinical samples (Figure 2C).

**Figure 2.**
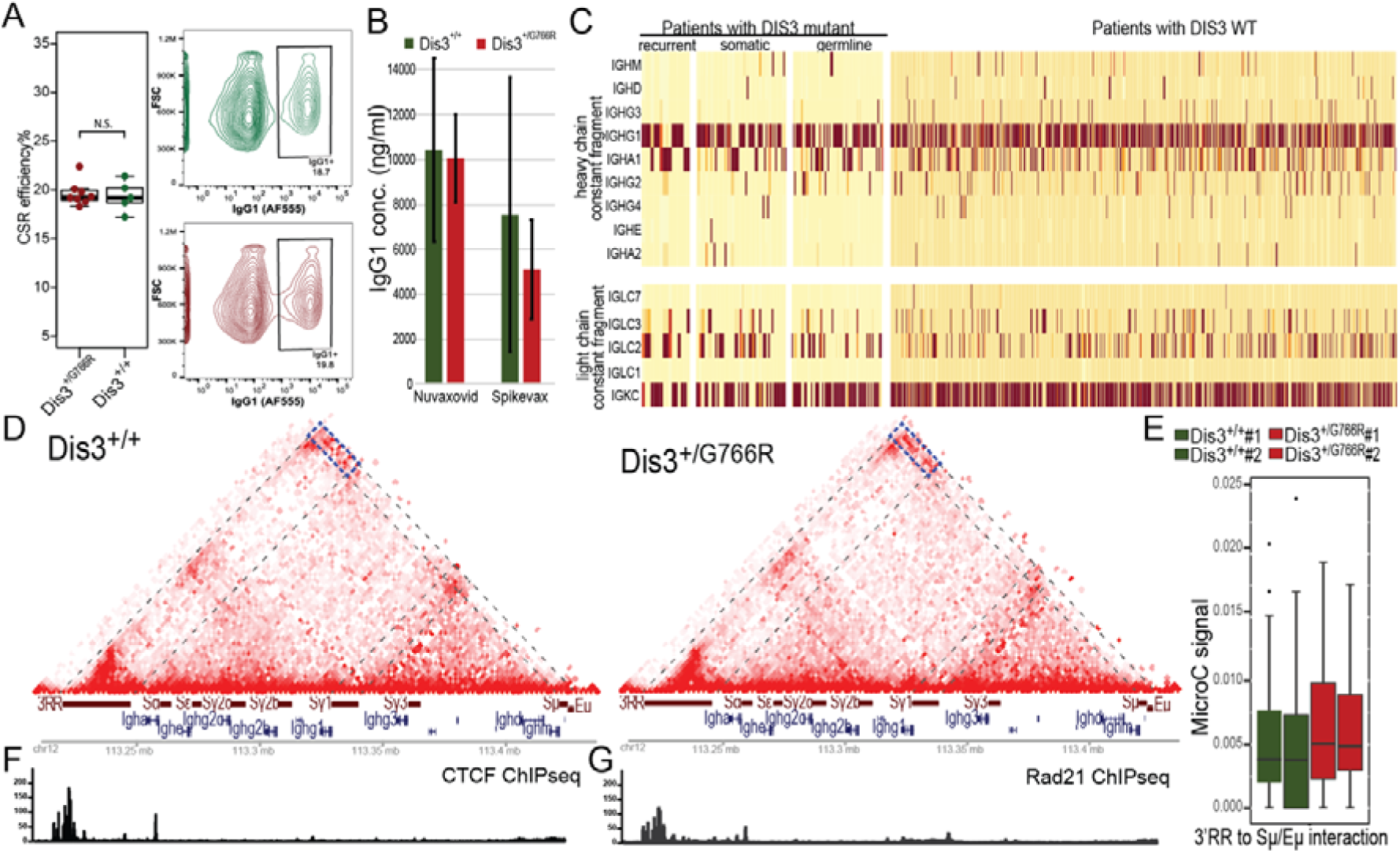
Unaffected immunoglobulin class switch recombination in B and MM cells with DIS3 MM variants (A) No significant changes in frequencies of switching to the IgG1 class are observed in WT and Dis3^G766R/+^ B-cells 3 days after activation with lipopolysaccharide and interleukin-4 (*p* = 1, odds ratio = 0.9). (B) ELISA of α-SPIKE IgG1 antibodies, 14 days after immunization of mice with Nuvaxovid and Spikevax vaccines. (C) No significant changes in frequencies of class switching in WT DIS3 patients or patients with MM-specific DIS3 mutations, both somatic and germline, as determined by the analysis of immunoglobulin heavy chain classes expression in RNA-seq experiments. (D) MicroC analysis of chromatin architecture at the *Igh* locus in wild type and *Dis3^G766R/+^*. Splenic B-cells were activated for 3 days *in vitro*. Displayed is the normalized interaction matrix at the resolution of 2kb. Ligation frequencies from two replicates per condition were summed up before the normalization. (E) Distribution of the normalized MicroC signal representing DNA interactions between the 3′RR super-enhancer and the Eµ intronic enhancer in *Dis3^G766R/+^* and *Dis3^+/+^* primary B-cells (separate for each individual, see box in D for the annotation of the interacting genomic intervals). Differences in all pairwise comparisons are not statistically significant. (F) CTCF and (G) Rad21 ChIP-seq signal In the *Igh* locus.

As a next step in *DIS3^G766R/+^* model characterization, we wanted to assess the structure of the *Igh* locus chromatin architecture, which plays a crucial role in CSR. As we wanted to enhance our power to detect loops, we employed MicroC, a high-throughput micrococcal nuclease (MNase)-based chromosome conformation capture technique^58^. MicroC libraries were generated using activated splenic primary B-cells, with two independent replicates representing individuals for each genotype. We sequenced a total of ∼1.2 billion (around 600 milion reads per condition) MicroC reads. As expected, we observed a plaid pattern of intrachromosomal contacts in both wild-type and mutant B cells. The overall interactomes were similar between conditions; we detected 20,603 and 20,279 loops in the wild-type and heterozygous mutant B cells, respectively (Table S10 and S11). Our analysis confirmed the presence of the previously described loops anchored by 3’RR/Eµ and Sγ1/Sµ, in *Dis3^G766R/+^* and *Dis3^+/+^* (Figure 2D-G). Loop anchors intersected ChIP-seq peaks of CTCF and Rad21^68^ (Figure 2F-G). Moreover, the normalized MicroC signal representing DNA interactions between the 3′RR and the Eµ (blue rectangle, Figure 2D) did not show statistically significant differences between Dis3^G766R/+^ and Dis3^+/+^ B-cells (Figure 2E). These results contrast the previously reported observation in *Dis3* KO B-cells, which showed abolished interactions^38^.

Taken together, by introducing Dis3^G766R^ mutation in a heterozygous setting, we recapitulate the clinical markup of MM cells. We show that Dis3^G766R/+^ B cells, despite a mild phenotype primarily related to non-coding transcription, feature no overt changes in physiology. This includes maintaining exosome function critical for antibody maturation, CSR, SHM, and preserving the *Igh* locus’s chromatin architecture.

### MM-specific *Dis3^G766R^* mutation drives murine plasmacytoma

To check the potential general predisposition of *Dis3^G766R/+^*mice to develop cancer, 33 individuals of *Dis3^G766R/+^* and 30 WT mice were aged under standard conditions for over two years. We observed no significant effect of the MM-associated *Dis3^G766R/+^* allele on survival (Figure 3A), nor any signs of accelerated cancer development. To test for an increased spontaneous transformation, we have also cultured MEF cells of both genotypes according to the 3T3 protocol, which models early cancer development and genetic stability^59^. *Dis3^G766R/+^* MEFs did not show an increased frequency of spontaneous immortalization (Figure S3A), arguing against a generalized tendency toward cancerous transformation in cells with MM-associated DIS3 point mutations.

**Figure 3.**
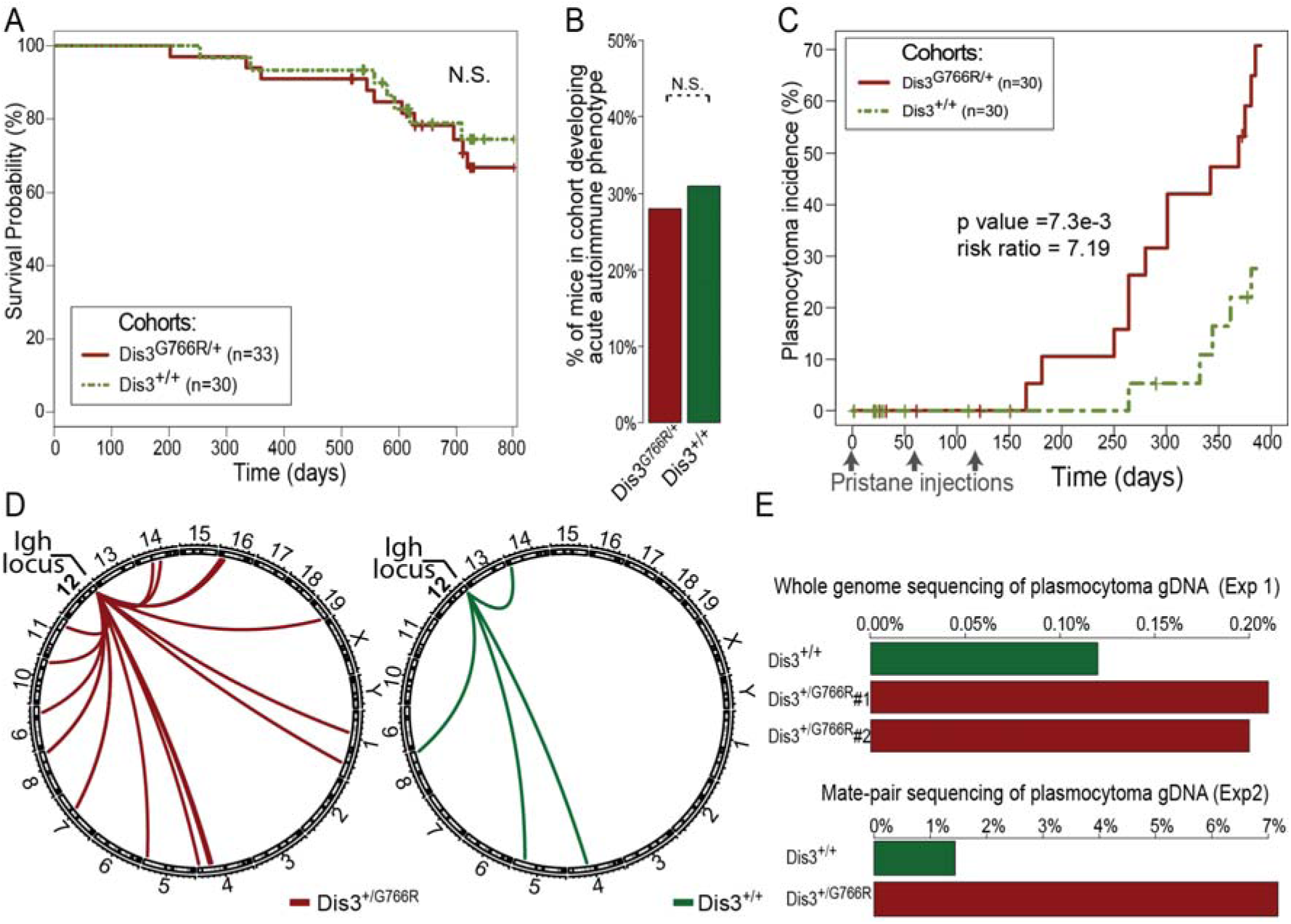
Predisposition to plasmacytoma development in Dis3^G766R/+^ knock-in mice is accompanied by frequent *Igh* translocations. (A) Kaplan-Meier survival plot of male WT and Dis3^G766R/+^ aged mice. (B) Incidence of WT and Dis3^G766R/+^ mice developing diffuse alveolar hemorrhage together with a generalized autoimmune phenotype in response to a peritoneal injection of pristane. (C) Cumulative incidence plot showing the frequency of plasmacytoma in Dis3^G766R/+^ mice and WT mice after the pristane injection. Note that the Dis3^G766R^ mouse line was constructed on a C57BL/6N genetic background, naturally resistant to plasmacytoma development. (D) Circos plot illustrates translocations that involve the *IGH* locus detected in plasmacytoma cells isolated from Dis3^G766R/+^ mice (red) and WT mice (green). (E) Fraction of chimeric reads in plasmacytoma samples isolated from Dis3^G766R/+^ and WT mice identified using standard whole-genome DNA sequencing (top) and the sequencing of mate-pare libraries (bottom).

Since *DIS3* mutations are specific to MM and not commonly found in other types of cancer, we decided to specifically investigate the development and progression of malignancies associated with a B-cell lineage. We employed a well-known *in vivo* model using pristane, an adjuvant that induces chronic inflammation, injection into the peritoneal cavity^13,14^, which leads to the development of plasmacytoma in susceptible mouse strains^60–62^. Both *Dis3^G766R/+^* and WT C57BL/6 mice exhibited similar immune responses to the pristane injection, indicated by the incidence of diffuse alveolar hemorrhage (Figure 3B). Despite the natural resistance of the C57BL/6 strain to plasmacytoma, the Dis3^G766R/+^ mice developed the disease, with a rate significantly higher than WT controls (risk ratio of 7.19; p = 0.0073; Figure 3C). Plasmacytoma cases displayed characteristic symptoms with atypical plasma cells in an ascites fluid (Figure S4B-D). Notably, *Dis3^G766R/+^* mice developed the condition not only more frequently and rapidly but also accumulated more ascite fluid than WT (Figure S4E-H).

As plasmacytoma is known to depend on aberrant AID-dependent translocations during CSR^13,14^, we wanted to investigate the frequency of translocations in Dis3^G766R/+^ and Dis3^+/+^ mice. DNA was isolated from the plasmocytomas, and deep sequencing was performed. The analysis of mate-pair and whole-genome sequencing data revealed that plasmacytomas from *Dis3^G766R/+^* mice had significantly higher rates of chromosomal translocations involving *Igh* locus (Figure 3D and Table S12 and S13). In addition, more chimeric reads originating from two different genomic locations were identified in *Dis3^G766R/+^*(Figure 3E), indicating increased genome structural instability. Because plasmacytomas are very rare in C57BL/6 mice and largely dependent on the Dis3^G766R/+^ genotype, additionally, we backcrossed Dis3G766R/+ mice to the plasmacytoma-prone BALB/c background, where plasmacytomas develop more frequently, providing sufficient WT control material for comparison. Notably, the difference in translocation-indicating reads between Dis3^G766R/+^ and WT genotypes was observed in both mouse strains (Figure 3E).

These results imply that the oncogenic effect of the *Dis3^G766R/+^* variant is exerted through the increased mutator effect driving translocations, particularly from the *Igh* locus, which agrees with a prevalence of *DIS3* mutation in non-HDR MM.

### Dis3-dependent translocations in mouse plasmacytomas have AID target characteristics

Considering that transient transcription and its degradation of the *Igh* switch locus is essential for AID-dependent recombination^20,39,45^, we aimed to characterize translocations in *Dis3^G766R/+^* plasmacytomas.

A meta-analysis of genomic regions surrounding chromosomal breaks detected in plasmacytomas revealed striking enrichment of AID occupancy determined previously by chromatin immunoprecipitation (ChIP-seq) (Figure 4A). Further analysis showed that both sites of genomic structural variants (SV) and regions of AID activity are GC-rich (Figure 4B), enriched with AID-specific motifs (Figure 4C), and are in close proximity to regions predicted to form G-quadruplexes (Figure 4D). RNA-seq read coverage over SV regions indicates enrichment of steady-state RNA levels over the random genome (Figure 4E-F, Figure S5). Most of them showed low-level pervasive transcription known to be predominantly cleared by DIS3. When transcribed (Figure 4G), regions of translocations are known to be prone to RNA polymerase stalling and R-loop formation. Indeed, regions involved in structural chromosomal aberrations in our plasmacytoma samples show a level of DRIP-seq accumulation (Figure 4H-J, Figure S5) like AID off-target loci (Figure 4H)^63^. Moreover, *Dis3^G766R/+^* B-cells accumulate and display an asymmetric distribution of RNA/DNA hybrids with a 5’ to 3’ slop on both DNA strands (Figure 4H). Since DIS3^G766R^ protein is less efficiently degrading structured RNA^18,19^, the observed asymmetry could reflect the staling of mutant Dis3 along its processive 3’ to 5’ activity. Analogously, the switch regions in the *Igh* locus show a roughly 25% increase in *Dis3^G766R/+^* DRIP-seq signal (Figure 4I). This can be explained by the fact that Dis3 mutants, including the MM *Dis3^G766R^* variant, are expected to have difficulties processing G-quadruplexes enriched in translocating regions (Figure 4D) and representing very stable RNA secondary structures. Similarly, the TSS regions of MM driver genes, known as AID targets^63^, have a significantly stronger DRIP-seq signal compared to the average of all active genes in B lymphocytes (Figure S5F). Consequentially, these regions in plasmacytoma are expected to be susceptible to AID-dependent DNA damage^46,64^. A snapshot of the immunoglobulin heavy chain, constant fragments genomic region (the best studied in the context of AID activity) exemplifies all the correlations mentioned above (Figure 4K).

**Figure 4.**
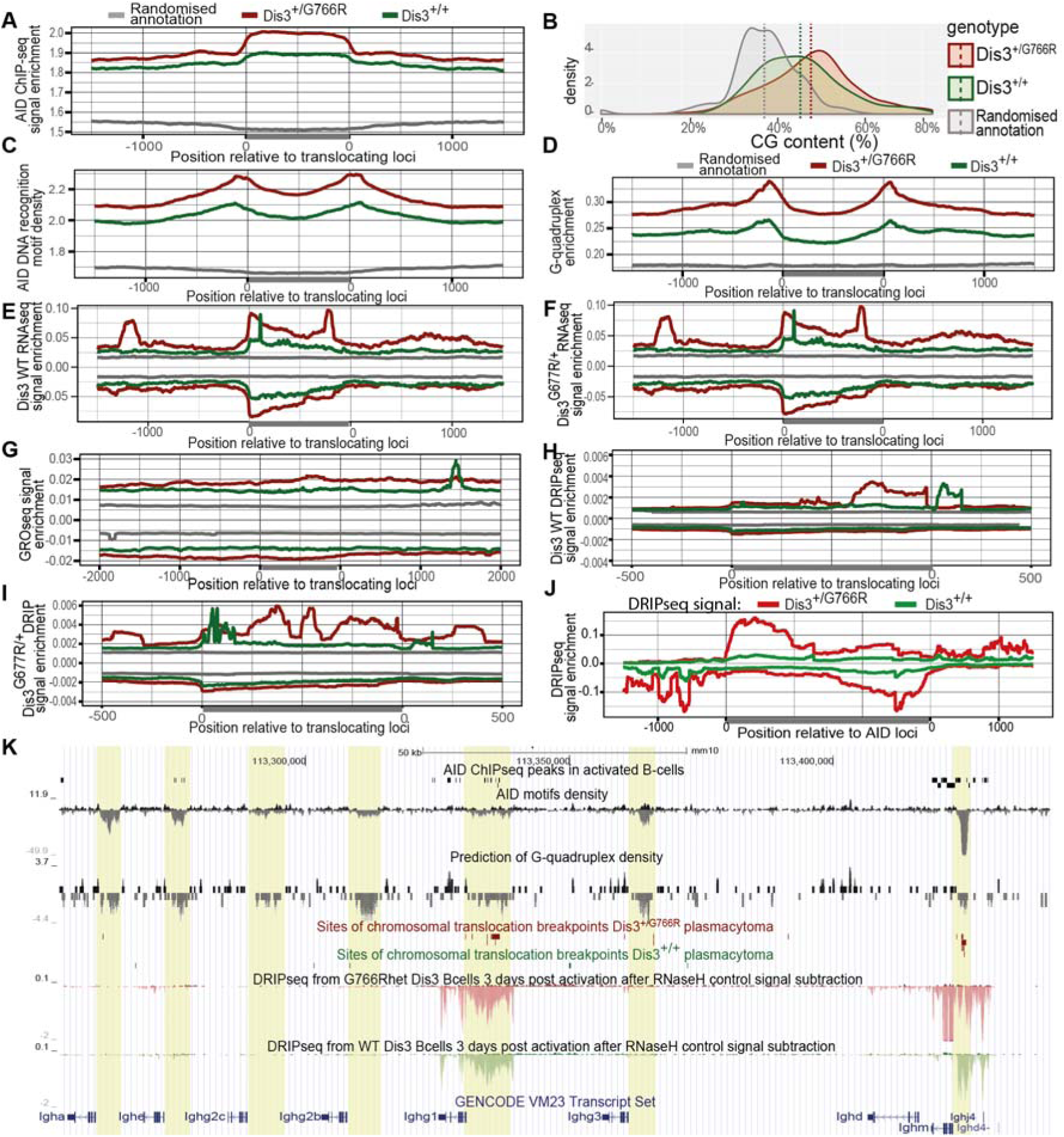
DIS3-dependent translocations in mouse plasmacytomas have AID target characteristics. (A-J) Meta-analyses of genomic regions engaged in translocations with features specific for sites of AID activity: AID ChIP-seq signal (A), GC-rich regions (B), AID-specific DNA recognition sequence motifs (C), predicted G-quadruplex sequences (D), WT (E) and Dis3^G766R/+^ (F) RNA-seq of day 3 activated B-cells, native transcription measured by GRO-seq (G) and WT (H) and Dis3^G766R/+^ (I) DRIP-seq signal of day 3 activated B-cells, relative to random genomic distribution. (J) Meta-analyses of DRIP-seq signal from Dis3^+/+^ and Dis3^G766R/+^ over known AID target loci. (K) UCSC genome browser snapshot of a fragment of the *IGH* locus that shows the clustering of translocation sites with regions that coincide with the enrichment of AID-specific recognition sequence motifs, G-quadruplexes, R-loops, active bidirectional transcription, and AID occupancy.

We conclude that chromosomal translocations caused by MM-associated DIS3^G766R^ mutation display AID target characteristics.

### Mutational profiles revealed signatures of promiscuous AID activity and a prevalence of translocations in patients with MM *DIS3* alleles

Having established that specifically in B-cell lineage MM *DIS3* alleles lead to increased genome instability, our objective was to test this finding against clinical data from the CoMMpass MM study, comparing patients with and without *DIS3* mutations. Patients were categorized based on the type of *Dis3* mutation, distinguishing between the recurrent mutations (D479, D488, and R780), other *DIS3* variants, and WT *DIS3*. These groups consisted of 22 patients with recurrent mutations, 48 patients with other variants, and 993 patients with WT DIS3, respectively.

First, analysis of translocations involving *IGH* locus revealed their significantly higher occurrence in patients with *DIS3* mutations (64.3% of non-recurrent *DIS3* mutated patients, 64.5 % of recurrent *DIS3* mutated patients, and 33.2 in the rest of the patients) (Figure 5A, B), agreeing with the specificity of *DIS3* mutations for non-HD MM and the role of *DIS3* in AID-dependent translocations.

**Figure 5.**
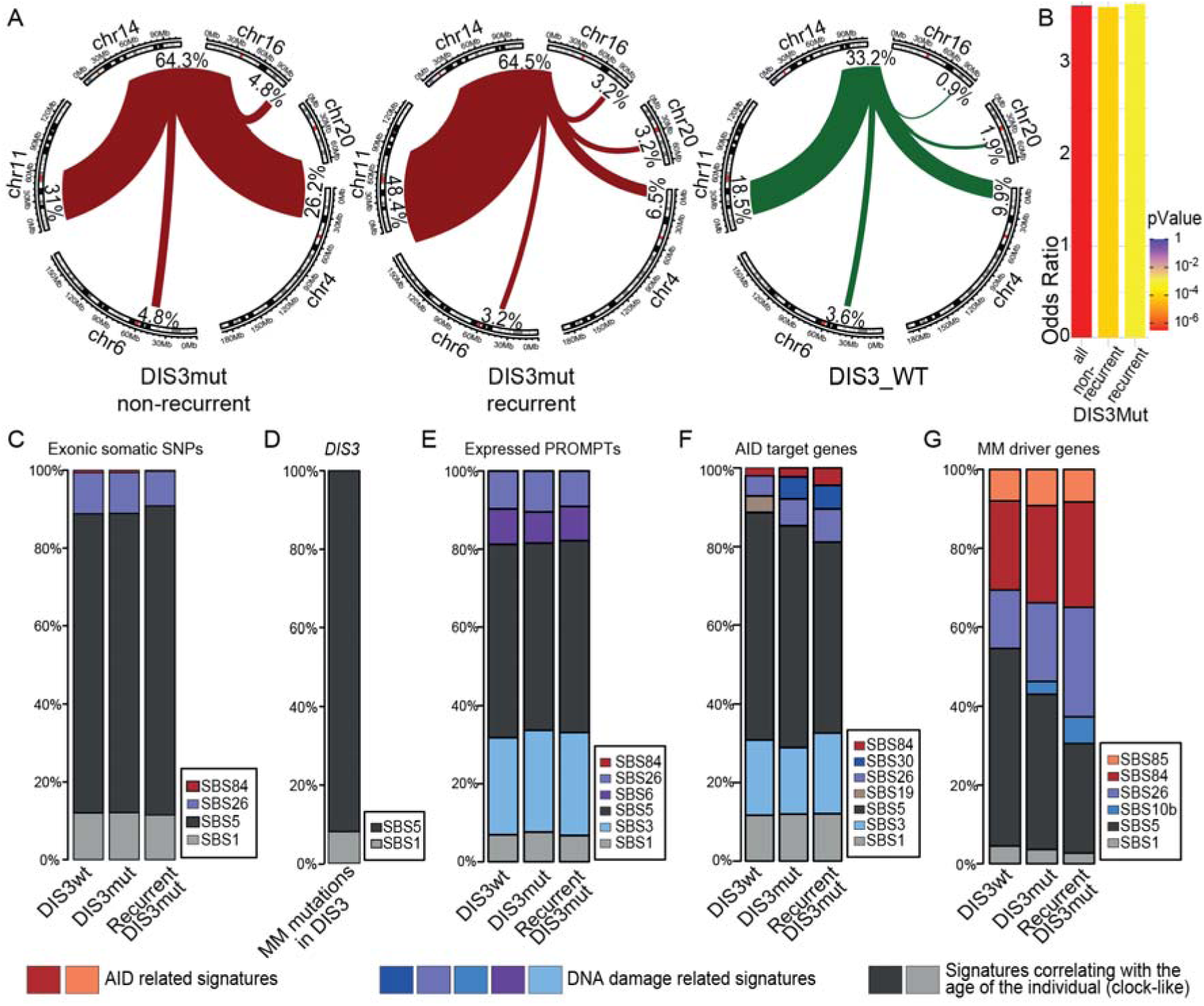
Mutational signatures reveal increases in AID and DNA damage repair lesions on MM driver genes in DIS3 mutant patients. (A) Circos graphs depicting the MM driver translocations of the *IGH* locus in WT DIS3 CoMMpass patients *vs*. patients with somatic non-recurrent and recurrent DIS3 MM variants. (B) Barplot of the enrichment of *IGH* MM-specific translocations in DIS3 mutant patients of different genotypes compared with WT DIS3 patients. (C-G) Contribution of COSMIC mutational signatures in all somatic single-nucleotide polymorphisms (SNPs) acquired by CoMMpass patients with WT DIS3, a mutant DIS3, or a recurrent MM DIS3 variant in whole exome sequencing (WES): exom-wide (C), in all somatic DIS3 mutations (D), PROMPTs in genes known to be AID off-targets (E), in genes known to be AID off-targets (F) and in MM driver genes (G).

Then, to characterize the effect of MM *DIS3* alleles on point mutations, we analyzed 96 COSMIC (Catalogue of Somatic Mutations in Cancer) mutational classes of single-base substitution (SBS) profiles^65^, estimating the contribution of each of them (Figure 5C-G, Figure S6C, D). We identified age-associated, exome-wide SBS5 as the primary mutational signature in patients of all *DIS3* genotypes (Figure 5C, Figure S6C), in agreement with the late onset of MM and previous analysis^15,66,67^. Similarly, MM-specific mutations detected in *DIS3* in CoMMpass study patients and mutations reported in the COSMIC database show a predominance of age-associated SBS5 (Figure 5D). Despite the exosome complex involvement in SHM, only a small fraction of mutations across the exome could be attributed to mutation signatures associated with AID activity across all the *DIS3* genotypes (Figure 5C), specifically SBS84 (reflecting canonical AID according to the most recent classification) and SBS85 (concordant with indirect effects of AID activity)^69^.

To be more specific, we next analyzed transcribed PROMPT regions toward which DIS3 is targeted. These indeed displayed distinct somatic mutation patterns compared to the entire exome. PROMPTs featured a significantly higher proportion of substitutions that can be attributed to DNA damage, especially SBS3, associated with the erroneous repair of double-strand DNA breaks in a BRCA-dependent mechanism (Figure 5E). This, in principle, could be explained by the reported DIS3-dependent increase in genome instability accompanying RNA:DNA hybrid accumulation^56^. However, despite PROMPTs being the most prominent DIS3 substrates, the somatic MM *DIS3* alleles do not affect their AID-depended mutagenesis (Figure 5E). Moreover, the contribution of AID-dependent signatures in all transcribed PROMPT regions was limited (Figure 5E).

To narrow down the search for the effect of DIS3 mutation even more, we analyzed known AID targets, which also include a subset of PROMPTs. There, the contribution of AID-dependent signatures is significant and higher in patients with MM *DIS3* alleles, especially the recurrent ones (Figure 5F). Further analysis of genomic regions encompassing known AID-dependent MM driver genes^63^ where mutations are under positive selection pressure during the carcinogenesis revealed a very strong overrepresentation of signatures related to both AID-dependent mutagenesis and defects in the repair of DNA lesions typical for CSR and SHM (namely: BER SBS10b, SBS30, and MMR SBS6, SBS26) (Figure 5F-G). Gratifyingly, patients with MM-specific DIS3 variants exhibit a drastic increase in the contribution of these signatures, especially SBS26 (35% in recurrent mutated DIS3 patients, 23% in non-recurrent mutated DIS3 patients, and 15% in the rest of the patients; Figure 5E). These results strongly support the positive effect of MM DIS3 variants on AID-dependent mutagenesis.

All analyses presented above indicate that although MM-associated DIS3 alleles do not alter the mutational profile genome-wide, they enhance AID-related mutagenesis occurring within a very limited time window of naive and memory B-cell activation rounds. Furthermore, these mutations are positively selected in very specific loci during MM development.

### Mutant DIS3 stalling on chromatin-bound substrates, resulting in increased DNA damage in activated B-cells

Given the correlation between MM *DIS3* alleles with AID-dependent mutations and translocations in both patients’ samples and the murine model, we next wanted to gain insights into the mechanism, which would have to be related to DIS3 chromatin association.

We found the immunoprecipitation of DIS3 on chromatin to be extremely challenging, possibly due to an indirect and transient nature of the DIS3 interaction with chromatin. Thus, to experimentally verify the direct involvement of MM DIS3 protein variants, we analyzed chromatin occupancy of the G766R mutant and WT DIS3 protein using the DamID methodology. This approach is based on fusing a protein of interest to DNA adenine methyltransferase (Dam), which modifies adjacent adenines in DNA, providing sufficient sensitivity to detect even weak interactions between DNA and its associated factors^70^. We modified the established mouse B-cell lymphoma CH12F3-2A cell line capable of cytokine-induced *Igh* CSR. Even before analyzing the sequencing data, we noticed 2.5-fold higher library yields from cells that express DIS3^G766R^ compared with WT DIS3 (Figure 6A). Such an increase can be explained by a higher chromatin occupancy of the mutated protein and agrees with the previously observed stalling of DIS3 mutant on RNA substrates^19^. DIS3 has a very broad pattern of association with chromatin. However, PROMPT regions, which give rise to the most potent DIS3 substrates (Figure 6B), and sites of translocations mapped in plasmacytomas show significant enrichment for DIS3 of both genotypes, coinciding with native transcription (Figure 6B,C). Notably, known AID off-target genes are among the most highly DIS3-occupied loci in the genome (Figure 6B). Moreover, DIS3 occupies the vicinity of switch regions in the *Igh* locus (Figure S6E), agreeing with the role of DIS3 in CSR.

**Figure 6.**
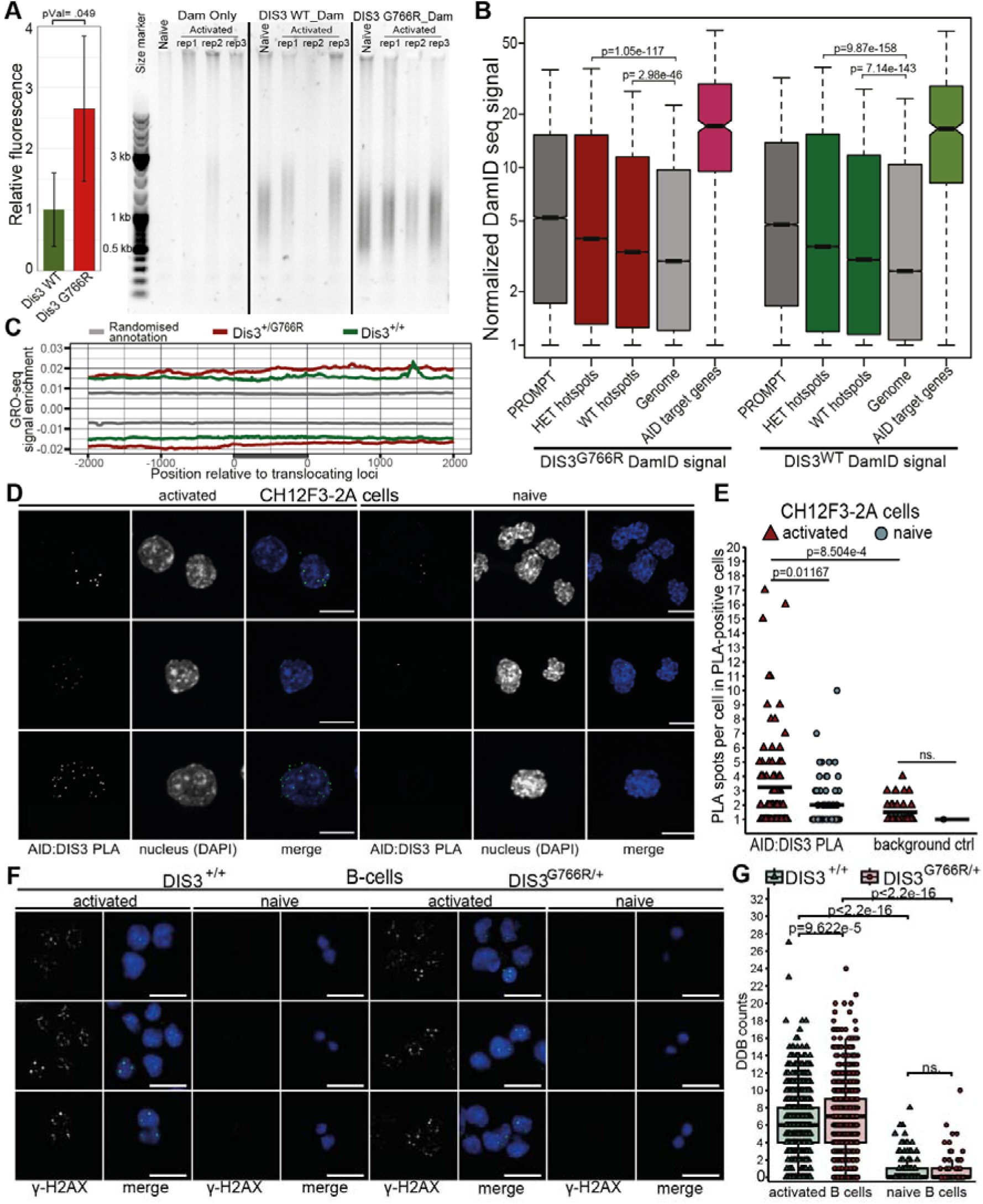
Mutant DIS3 stalling on chromatin-bound substrates, resulting in increased DNA damage in activated B-cells. (A) Quantification of intensities of DamID libraries obtained from activated CH12F3-2A cells that express WT and DIS3^G766R^. Error bars represent the standard deviation. (B) DIS3 DamID-seq in activated CH12F3-2A cells shows signal enrichment compared with the rest of the genome over regions that translocate in plasmacytomas and represent known DIS3 substrates, such us PROMPTs (Mood’s median test). (C) GRO-seq shows the enrichment of native transcription at the same genomic sites. (D) Representative confocal microscopy images demonstrating DIS3 and AID interaction in CH12F3-2A cells detected by *in situ* PLA (interaction site in green, merged with DAPI in blue). Only the negative control giving the highest background among is shown. (E) Quantification the number of DIS3 and AID interaction sites (PLA spots) in CH12F3-2A. Analysis was restricted to cells with at least one PLA spot identified (PLA-positive cells). Statistics were performed with the Wilcoxon test. The black horizontal lines represent means. (F) Representative images demonstrating γ-H2AX staining in naïve and day 3 activated B-cells (green, merged with DAPI in blue). (G) Quantification of number of γ-H2AX immunofluorescent spots per single cell. Statistics were performed with the Wilcoxon test. Scale bars, 10 μm.

The data presented above indicate that DIS3 acts together with AID to drive translocations. To support cooperation with an orthogonal methodology, we have analyzed interactions between DIS3 and AID using Proximity ligation assay (PLA) (Figure 6D). Indeed, in CH12F3-2A cells, we see PLA-positive spots, which number is higher upon *in vitro* activation (Figure 6E and Figure S6F). We also examined DNA damage accompanying B-cell activation *in vitro* using γ-H2AX, an established marker dsDNA breaks^71^. Naïve B-cells present a low level of DNA damage, which, however, increased significantly in cells 3 days post-activation (Figure 6F,G). In contrast to naïve, activated B-cells demonstrated a moderate, yet statistically significant, increase of DNA damage in *Dis3^G766R/+^* compared to *Dis3^+/+^* (Figure 6F,G), further supporting the role of MM *DIS3* alleles in B-cell specific cancerogenesis.

These results imply that the oncogenic effect of *Dis3* mutations is exerted through the direct impact of the mutant protein on DNA rather than by the accumulation of its RNA substrates.

### *Dis3^G766R^* allele does not affect the chromatin architecture but hijacks it for joining enhancers with proto-oncogenes

DIS3 and the RNA exosome complex were postulated to be essential for suppressing eRNAs from super-enhancers^45^ and PROMPTs at active TSSs^72^. We therefore tested whether translocations in plasmacytomas correlate with these regulatory regions. Using publicly available ChIP-seq data from B-cells 3 days post-activation^68^, we defined active enhancers and promoters as regions significantly enriched for H3K4me1 and H3K4me3, respectively (FDR < 0.01), overlapping with H3K27ac peaks. Translocating loci were localized in vicinity to either enhancer-specific H3K4me1 enrichments or active promoter-specific H3K4me3 ChIP-seq peaks with a median distance of 33kb or 25kb for loci detected in WT and *Dis3^G766R/+^* plasmacytomas, respectively (Figure 7A and S6G). The annotated chromosomal breaks also exhibited a non-random distribution with respect to their proximity to p300 ChIP-seq peaks (Figure 7B and S6I) a marker of active enhancers as well as super enhancers (Figure S6H).

**Figure 7.**
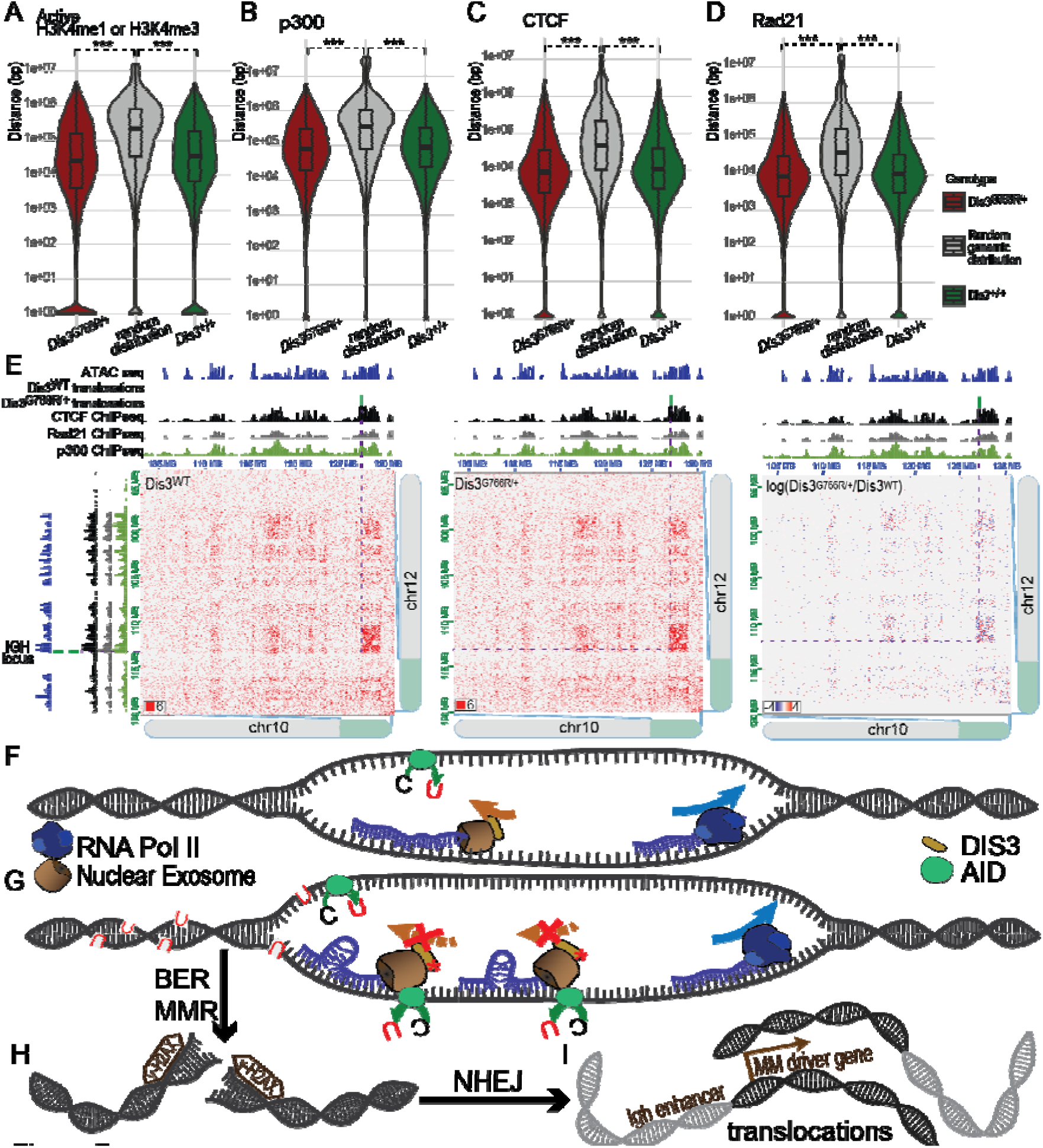
Model of the involvement of MM-specific DIS3 variants in MM oncogenesis through enhanced AID recruitment to ssDNA. (A-D) Distributions of distances of regions translocating in murine plasmacytomas to the closest ChIP-seq peaks of: H3K4me1 or H3K4me3, both overlapping with H3K27ac (A), p300 (B) CTCF (C) and Rad21 (D). Statistics were performed with the Wilcoxon test. (E) Representative example of an Intrachromosomal interaction accompanying Igh-derived translocation t(10,12) in murine-induced plasmacytoma. (F) AID is recruited to DNA during class switch recombination (CSR) through transcription-generated single-stranded DNA regions. These regions serve as substrates for AID, initiating CSR and ultimately producing antibodies with altered functions. The nuclear exosome with WT DIS3 recruits AID to transcribed sequences, but due to the transient AID activity outside the *Igh* locus is a rare event. (G) MM DIS3 variants increase AID activity by enhancing the nuclear RNA exosome affinity, resulting in increased occupancy/stalling. This results in the accumulation of uracil bases in off-target regions. (H) DNA damage repair mechanisms, including base excision repair (BER) and mismatch repair (MMR) involving uracil glycosylase and AP-endonucleases, process AID introduced uracils, inducing double-strand breaks (DSBs). (I) Non-homologous end joining (NHEJ) rejoins the free DNA ends, but excessive DSBs can lead to unrestrained genome rearrangements.

To further investigate the relationship between chromatin architecture and translocations, we studied TADs by analyzing positions of CTCF and Rad21 ChIP-seq peaks identified in B-cells 3 days post-activation^68^ in relation to the translocations found by us in plasmacytomas. Indeed, translocating loci were localized significantly closer to CTCF and Rad21 ChIP-seq peaks than expected from a random distribution in the genome (Figure 7C,D, and S6J). We detected a median distance of translocating loci from the CTCF peak to be 10.1kb or 8.6kb, and from the Rad21 peak to be 10kb or 8.5kb, in WT and *Dis3^G766R/+^*plasmacytomas, respectively.

Since, in contrast to *Dis3* ablation, the *Dis3^G766R^*allele does not interfere with the chromatin architecture of the *Igh* locus, we closely examined MicroC data for translocations originating in the *Igh* locus. Translocations identified in the plasmacytomas are in regions with strong interaction, indicating that they are in close physical proximity within the three-dimensional nuclear space (Figure 7E and S7). Similarly to the *Igh* locus, these interactions do not depend on the Dis3 variant (Figure 7E and S7).

In sum, we demonstrated that although the Dis3 mutant does not alter chromatin architecture, it increases the frequency of translocations between genomic regions involved in transcription during B-cell activation. This leads to a permanent joining of active promoters and enhancers in developing B cells.

## DISCUSSION

A common property of cancer cells is alteration of gene expression. This was initially attributed mainly to changes in transcriptional regulation but more recently also to posttranscriptional regulatory mechanisms. In hematologic malignancies, mutations of spliceosome components, such as SF3b^73–76^ and U2AF^73,77^, affect alternative splicing patterns. Still, mechanistic links are missing despite many clinical and functional analyses employing several transgenic mouse models that show effects on RNA metabolism^78^. In MM, two genes involved in RNA metabolism DIS3^17^ and TENT5C (FAM46C)^79^ are specifically mutated^7,8,16^, jointly being affected in almost 30% of MM patients. TENT5C is a MM specific oncosupressor^7,8,79^. In normal B-cells, TENT5C enhances the expression of immunoglobulins by stabilizing their mRNA through cytoplasmic polyadenylation, which at the same time increases the protein load into the endoplasmic reticulum^80^. TENT5C disruption decreases ER stress and enhances proliferation in antibody-producing malignant plasma cells.

Our present study provides a mechanism by which DIS3 drives oncogenic transformation in MM. Contrary to previous suggestions, dysregulation of the transcriptome is not the cause of the neoplastic transformation. MM *DIS3* alleles also do not lead to ubiquitous genome destabilization. Instead, MM-associated *DIS3* mutations lead to a gain-of-function, which introduces B-cell-specific destabilization of the genome, enabling the acquisition of genetic lesions and giving rise to oncogenic genome rearrangements, such as *IGH* translocations, known to be the primary drivers of MM (Figure 7F).

DIS3 is a primary nuclear exoribonuclease that acts from the RNA 3’ end. It targets many types of RNA species, mainly pervasive transcription products that are rapidly cleared. In a mouse model, a relatively mild heterozygous *Dis3^G766R^*, the accumulation of the most potent DIS3 substrates is focal, localized mainly to unannotated, non-coding regions of the genome, and shows no detectable changes in the protein-coding transcriptome as well as the proteome. This can be attributed to the fact that DIS3^G766R^ partially retains some of its activity showing a significant inhibition over some substrates, specifically structured RNA^18^. DIS3^G766R^ stalling on structured RNA substrates results in increased affinity of the mutant DIS3 to chromatin. In B cells, DIS3 is involved in SHM and CSR, processes that require the activity of AID. Chromatin-stalled DIS3 variants associated with multiple myeloma (MM) facilitate the recruitment of AID, leading to increased and more promiscuous AID activity beyond its usual targets. This heightened activity results in a greater frequency of translocations, contributing to genomic instability and potentially driving oncogenic events (Figure 7F). In a mouse model, the heterozygous *Dis3^G766R^*^/+^ mutation increases the rate of plasmacytoma development, accompanied by a significantly higher level of structural genomic variants.

In physiological situations, there are activation-dependent interactions between DIS3 and AID in the B-cell lineage confirmed by us and restricted to limited spots, comparable to the numbers of DNA damage foci measured by γ-H2AX B-cell activation *in vitro*. The statistically significant but moderate increase of DNA damage in *Dis3^G766R/+^* cells is in contrast to *Dis3* depletions, which dramatically affect both cell physiology and genome integrity^56^. Moreover, MM *DIS3* mutations display no CSR impairments both in mouse models and in MM samples. The chromatin architecture of the *Igh* topologically associating domain, as well as the rest of the genome, does not show statistically significant changes. Both contrast with *Dis3* inactivation, which leads to the rearrangement of tertiary chromatin architecture and inhibits CSR in murine B-cells^38^.

The nuclear exosome is known to be involved in super-enhancer regulation by controlling levels of eRNA^45^. Interestingly, AID has been shown to promote off-target mutations and genomic rearrangements at super-enhancers, which contributes to oncogene activation and tumor progression^53^. Our data show that translocations detected in murine plasmacytoma predominantly originate in the proximity of enhancer regions as well as active promoters. Translocations originating from the *Igh* locus in plasmacytoma correlate with interchromosomal interactions identified through MicroC data. This aligns with existing evidence that the Igh locus participates in tertiary chromatin structures, interacting in a developmentally regulated manner with specific B cell lineage genes, such as Pax5, Ebf1, Aff3, and Foxp1, many of which overlap with loci implicated in translocations in early B cell malignancies^81^. MM-associated DIS3 mutations may exploit these pre-existing chromosomal interactions, enhancing the likelihood of aberrant translocations. Given the nuclear exosome’s role in RNA surveillance within regulatory elements of active gene networks, such mutations can hijack existing chromosomal tertiary structures. In such a way, MM-associated DIS3 mutations, even if they are biochemically subtle and have limited transcriptomic impact, can lead to significant consequences for oncogenesis. A restricted spectrum of DIS3-dependent mutagenesis in MM is supported by the analysis of mutational signatures in MM samples, clearly showing the association of *DIS3* mutations with AID-related lesions in MM driver genes and not genome-wide, which can occur only in a short window of opportunity during B-cell activation.

Interestingly, in MM patients, beyond the overall increase in driver translocations, there is a striking specificity for specific translocations depending on the *DIS3* genotype (Figure 5A,B). The t(4;14) translocation is depleted in patients with recurrent *DIS3* somatic mutations. At the same time, in patients with the coexistence of both lesions, the recurrent MM DIS3 variants always represent minor clones only. Notably, the t(4,14) translocation activating pro-proliferative FGFR3 expression in 90% of patients is accompanied by the del(13q) deletion encompassing the *DIS3* gene^82,83^. Thus, its coexistence with the defective MM-specific *DIS3* allele may lead to the faster and more effective counter-selection of recurrent *DIS3* mutation in patients with the t(4,14) translocation.

In summary, our study shows that mutations in the ubiquitously expressed RNA-degrading enzyme DIS3 enhance its chromatin affinity and AID-dependent mutator activity, driving cancer specifically in the B-cell lineage and contributing to multiple myeloma (MM) development. It clarifies the role of DIS3 mutations in MM development and explains the specificity of mutant DIS3-induced genetic lesions for MM or, in the case of the mouse model, plasmacytoma.

## Supporting information

Methods section

## Acknowledgments

We thank Justyna Chlebowska, Jakub Gruchota, Kamila Kłosowska-Kosicka, Dorota Adamska, Michał Kamiński, Radosław Salamon, Katarzyna Prokop, and Monika Kusio-Kobiałka for help with selected experiments, Maria Anna Ciemerych-Litwinienko, Dominika Nowis, Jakub Gołąb, Joanna Kufel, Katarzyna Matylla-Kulińska for attentive readings of the manuscript and all Andrzej Dziembowski lab members for fruitful discussions. We thank the expert support of the Mouse Clinic Institute (Illkirch) in mouse construction and handling. Efforts of the Multiple Myeloma Research Foundation (MMRF) and centers that contributed to the CoMMpass study are also acknowledged.

All mice lines were genotyped by the Genome Engineering Facility (GEF), part of IIMCB IN-MOL-CELL Infrastructure (RRID: SCR_021630) funded by the European Union – NextGenerationEU under National Recovery and Resilience Plan, Horizon Europe (Project 101059801 - RACE) and RACE-PRIME project carried out within the IRAP programme of the Foundation for Polish Science co-financed by the European Union under the European Funds for Smart Economy 2021-2027 (FENG) IIMCB.

This work was mainly supported by grant funding from the National Science Centre (NCN) to AD (UMO-2016/22/A/NZ4/00380; UMO-2013/10/M/NZ4/00299) and TK (UMO-2019/32/C/NZ2/00558), and grant funding from Ligue Contre le Cancer (Equipe Labellisée 2020), USIAS 2015, within the framework of ANR-10-IDEX-0002-02, ANR-10-LABX-0030-INRT, and ANR-17-EURE-0023 to BS. This research was co-supported by funding from the European Union’s Horizon 2020 research and innovation program (grant agreement no. 810425). Work in the Pękowska laboratory is funded by the Dioscuri Grant [Dioscuri is a program initiated by the Max Planck Society (MPG), jointly managed with the National Science Centre in Poland (NCN), and mutually funded by the Polish Ministry of Education and Science and the German Federal Ministry of Education and Research (BMBF)]; by the OPUS17 (UMO-2019/33/B/NZ2/02437), OPUS22 (UMO-2021/43/B/NZ2/02934), and Sonata Bis 11 (UMO-2021/42/E/NZ2/00392) grants from the NCN; and by the EMBO Installation Grant.

## Author contributions

TK: conceptualization, all bioinformatic analysis, and experimental work; OG, MM, JD, KK, ASC, MJS, EPO, MN: experimental work; AP, DC: bioinformatic analysis BS: conceptualization and knock-in mice; AD: conceptualization, supervision, original draft preparation.

