## Supplemental figures for "Multiple Myeloma associated DIS3 mutations drive AID-dependent IGH Translocations"

**Kuliński et al.**

### SUPPLEMENTAL DATA

- **Figure S1**
- **Figure S2**
- **Figure S3**
- **Figure S4**
- **Figure S5**
- **Figure S6**
- **Figure S7**

- **Description of Supplementary Tables**
- **Data, code and materials availability**

### Supplementary Figures

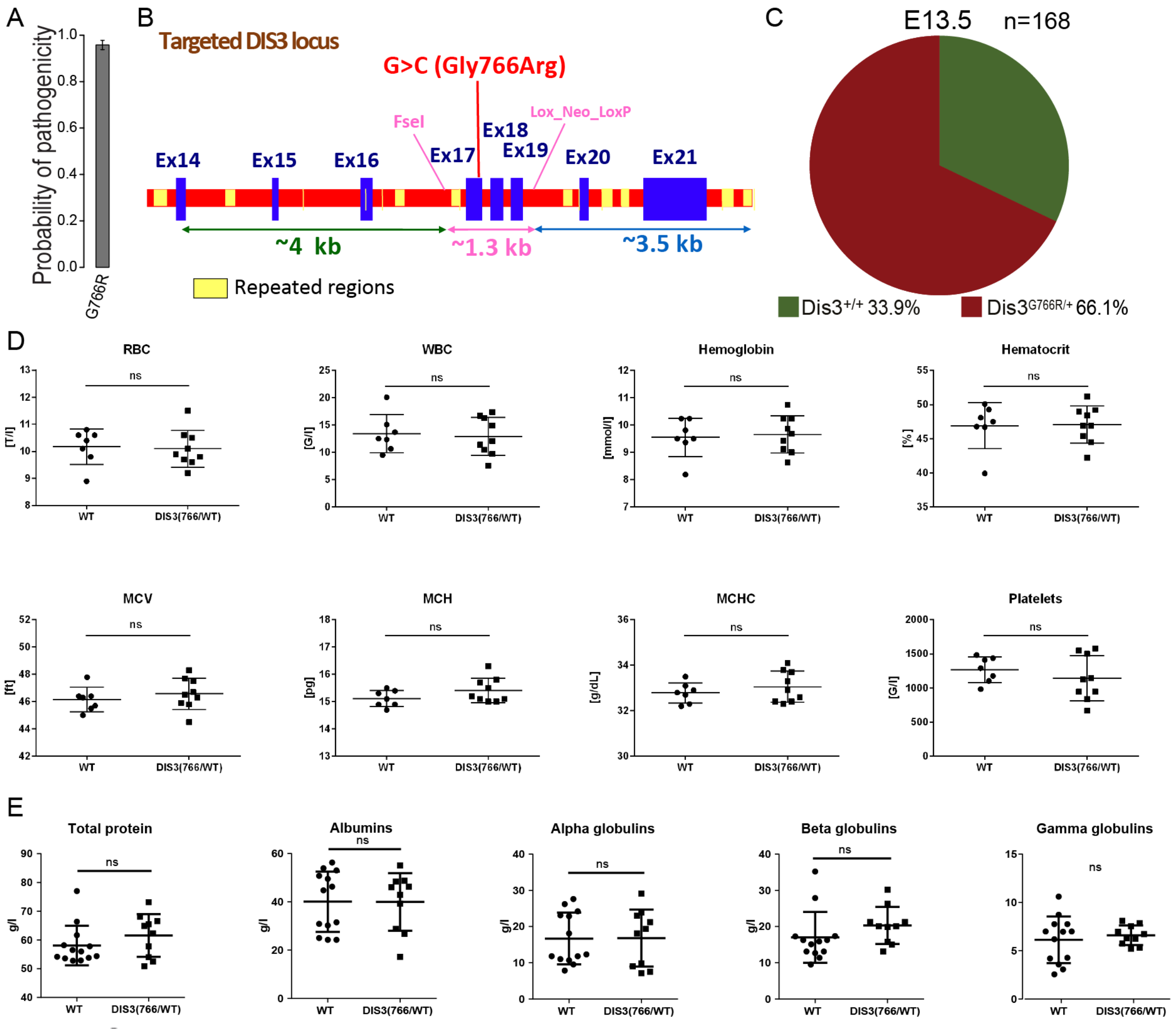


**Figure S1.** DIS3$^{G766R}$ allele leads to early embryonic lethality in homozygous mice, but heterozygous mice do not display any obvious abnormalities. (A) Prediction of G766R Dis3 variant pathogenicity by PON-P2. Error bars represent standard deviation. (B) Strategy for the generation of Dis3$^{G766R}$ knock-in mice. (C) Frequencies of offspring of Dis3$^{G766R/+}$ × Dis3$^{G766R/+}$ mattings on mouse embryonic day 13.5 (E13.5) of development with indicated genotypes. Dis3$^{G766R/+}$ mice do not show any abnormalities in (D) blood morphology or (E) disruptions of protein fractions in serum protein electrophoresis.

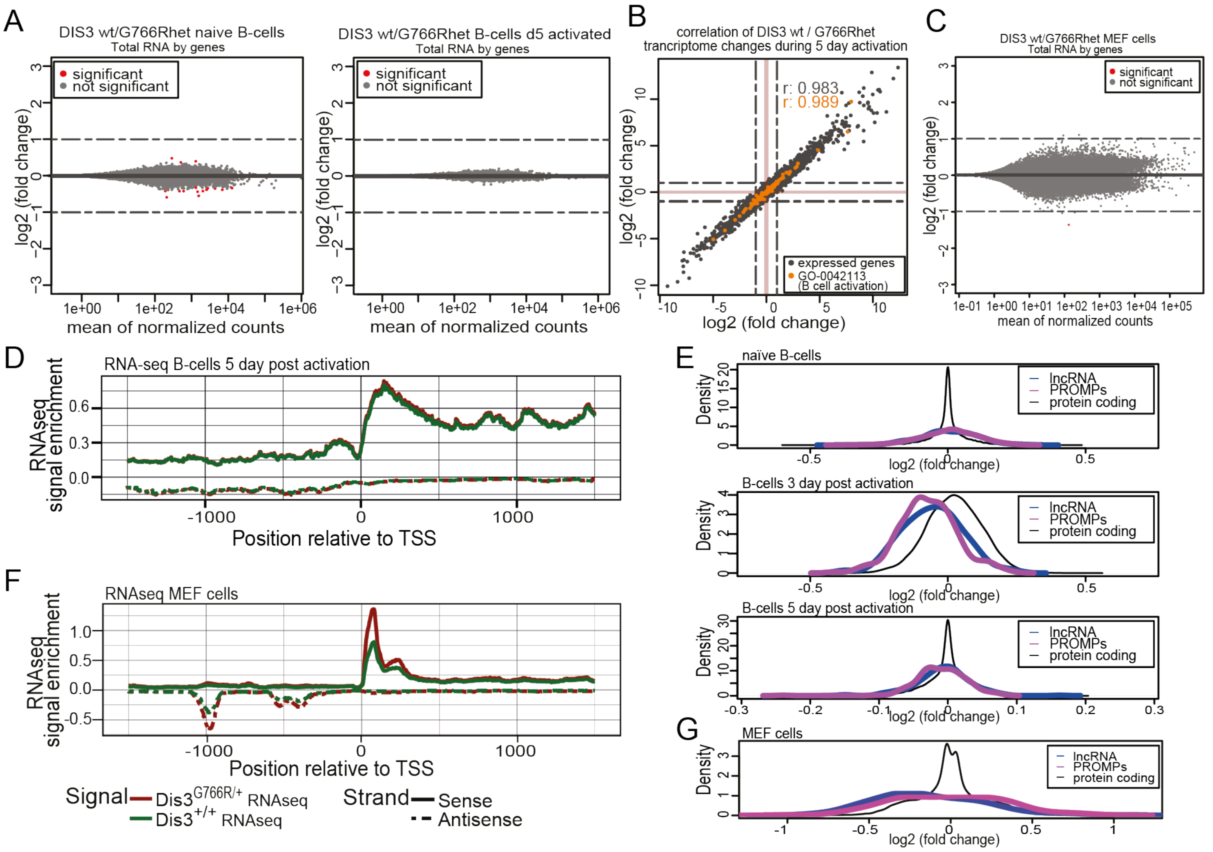


**Figure S2.** In vitro activated Dis3$^{G766R/+}$ B-cells present a molecular phenotype compared to WT B-cells. (A) MA plot comparing the transcriptome of primary Dis3$^{G766R/+}$ and Dis3$^{+/+}$ B cells naïve and *in vitro* activated for 5 days (n=3 for each genotype). (B) Correlation of transcriptome changes in the DIS3$^{G766R/+}$ and DIS3$^{+/+}$ during *in vitro* activation for 5 days. (C) MA plot showing comparisons of the transcriptome of WT MEFs with Dis3G766R/+ MEFs. The dashed lines on the plot represent a threshold of 2-fold change. (D) Meta-analyses of RNA seq signal over TSS of expressed genes day 5 activated B cells (E) The accumulation of PROMPT and long non-coding transcripts (lncRNA) in naïve, day 3, and day 5 in vitro activated B-cells. (F) Meta-analyses of RNA seq signal over TSS of expressed genes in MEF cells. (G) The distribution of log2 of fold change values shows a moderate accumulation of long non-coding transcripts (lncRNA) in DIS3$^{G766R/+}$ MEF cells, peaking at around 25% (~-1.25 fold change).

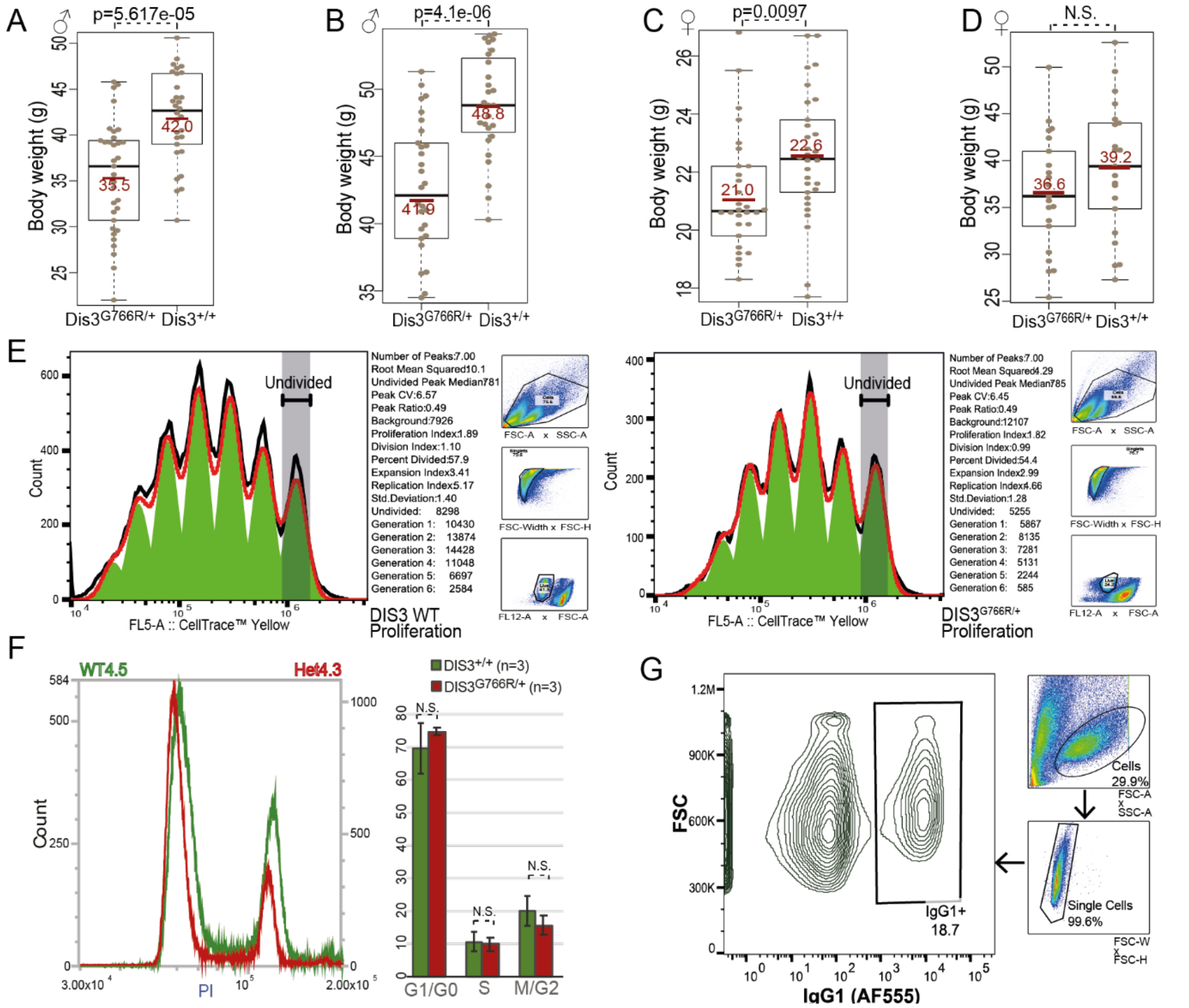


**Figure S3.** Generalized growth phenotype of Dis3$^{G766R/+}$ knock-in mice. (A-D) Weight of 25-45 week male (A), 60-80 week male (B), 16-25 week female (C), and 25-40 week female (D) Dis3$^{G766R/+}$ mice compared with their Dis3$^{+/+}$ littermates. (E) FlowJo proliferation tools analysis of representative Dis3$^{+/+}$ and DIS3$^{G766R/+}$ day 3 in vitro activated B-cell samples stained with CellTrace™ Yellow Cell proliferation assay. (F) Flow cytometric analysis of the cell cycle phase distribution estimating DNA content of MEF cells isolated from Dis3G766R/+ mice compared with their Dis3$^{+/+}$ littermates after 5 passages in cell culture, using propidium iodide (PI) staining. Quantification of mitotic phases in cells isolated from 3 litters of mice. (G) Representative flow cytometry analysis of primary activated B-cells that exemplify the gating strategy for the CSR assay

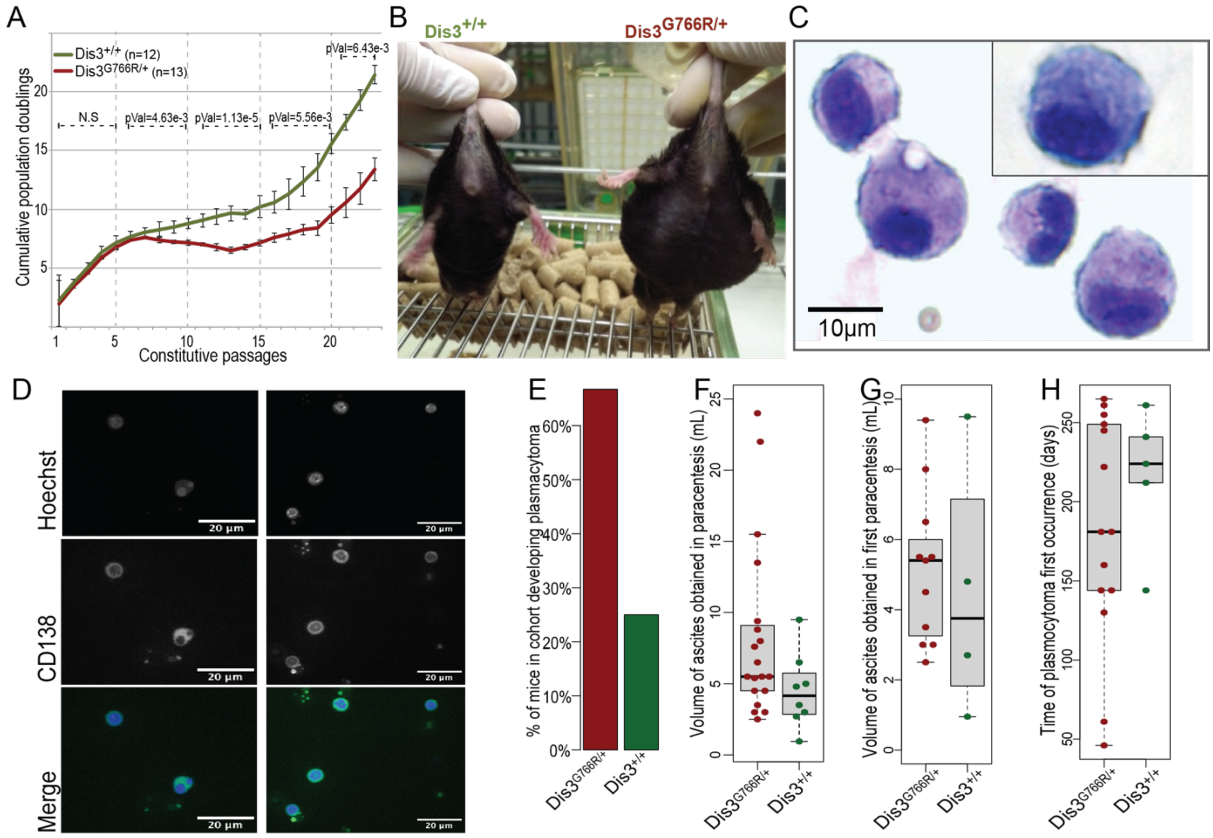


**Figure S4.** Characterization of plasmacytomas that develop in Dis3$^{G766R/+}$ and WT mice. (A) Cumulative population doubling of WT and Dis3G766R/+ MEFs cultured according to the 3T3 protocol. (B) Example of Dis3$^{G766R/+}$ mouse that developed plasmacytoma compared with its Dis3$^{+/+}$ littermate. (C) Representative smear of ascites fluid, showing atypical plasma cells. (D) Representative immunofluorescent staining with $\alpha$-CD138 antibody of atypical plasma cells smear in the ascites fluid. (E) Frequencies of plasmacytoma development in Dis3$^{G766R/+}$ mice and WT mice. (E) The volume (ml) of ascites that were collected during all paracentesis in Dis3$^{G766R/+}$ and Dis3$^{+/+}$ mice. (G) Volume (ml) of ascites that were collected during the first paracentesis in Dis3$^{G766R/+}$ and Dis3$^{+/+}$ mice. (H) Time of first plasmacytoma occurrence in mice that were treated with pristane.

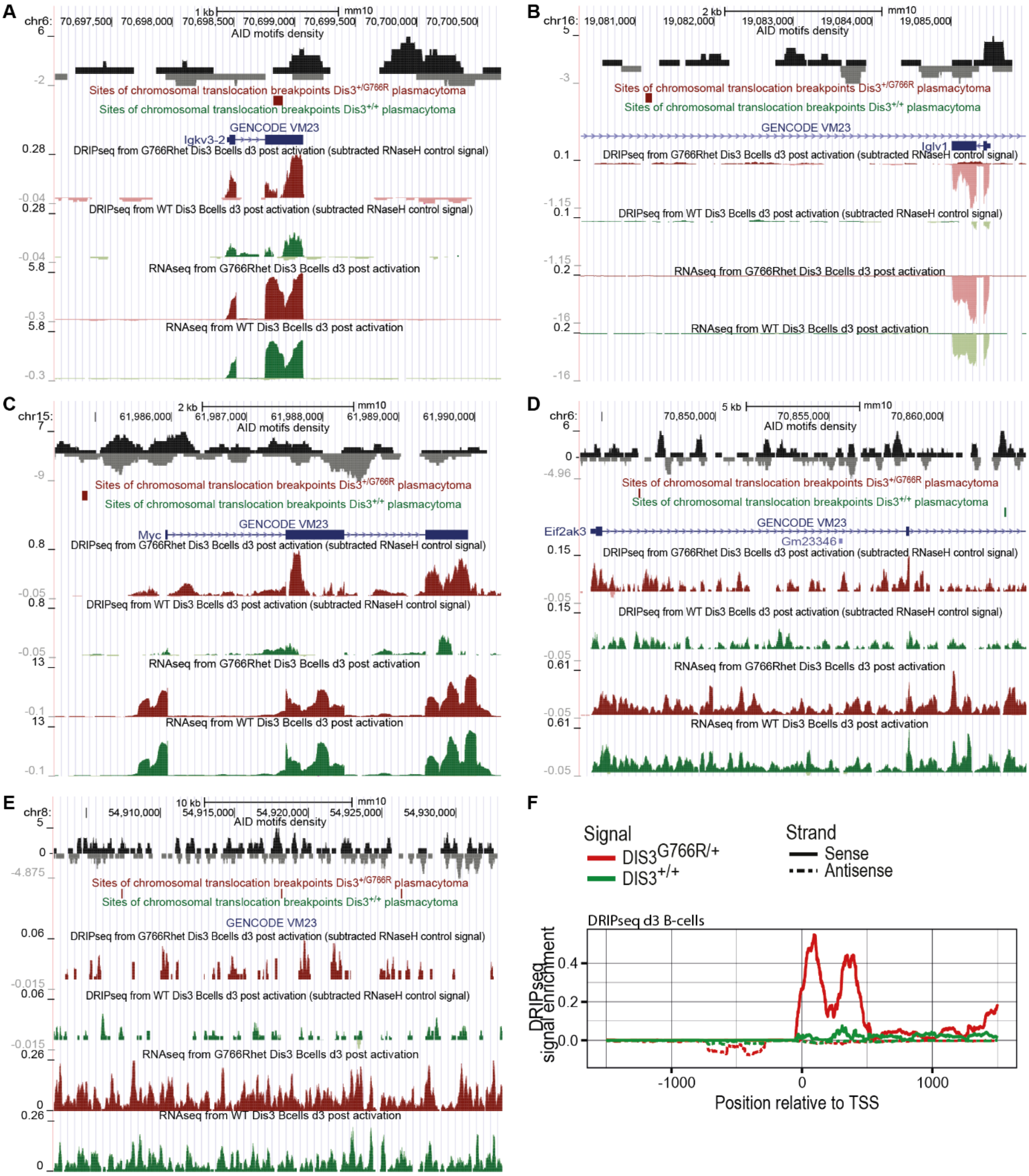


**Figure S5.** Representative translocating genomic regions in murine plasmacytoma: (A) locus of immunoglobulin light chain kappa variable region; (B) locus of immunoglobulin light chain lambda variable region; (C) *Myc* locus; (D) *Ei2fak3* locus with high level intronic transcription; (E) a locus of unannotated intergenic transcription on chromosome 8 a with 3 inter-chromosomal breakpoints. DRIPseq signal is calculated by subtracting the corresponding RNaseH background control. (F) Meta-analyses of DRIP-seq signal over TSS of MM driver genes, in day 3 activated B-cells.

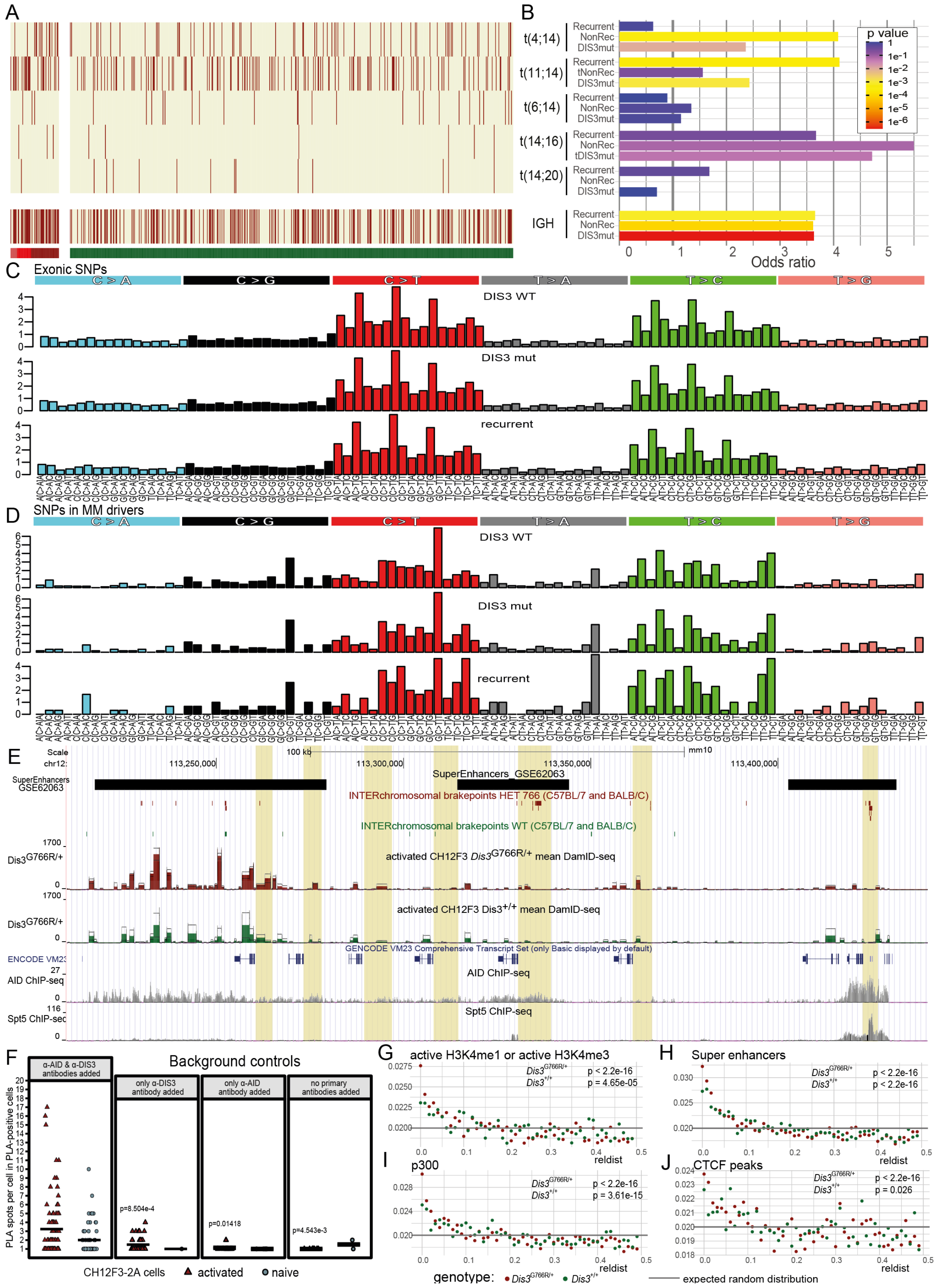


**Figure S6.** Mutational footprint of DIS3 variants in MM patients. (A) Heatmap of MM driver translocations of the IGH locus in WT DIS3 CoMMpass patients *vs*. patients with a DIS3 mutation and their subset with recurrent DIS3 MM variant. (B) Barplot of the enrichment of the IGH MM-specific translocations in the groups' odds ratios of translocation occurrence in DIS3 mutant and WT patients. (C) Mutational profiles of SNPs that were identified in whole exome sequencing (WES). (D) Mutational profiles of SNPs that were identified in WES in regions of known MM driver genes and their corresponding PROMPTs. (E) UCSC genome browser snapshot of the mouse IGH locus that shows $DIS3^{G766R}$ and $DIS3^{WT}$ occupancy over and in close vicinity of the switch regions, coinciding with AID and Spt5 occupancy and translocations that occur in plasmacytomas. (F)Quantification of *in situ* PLA detection of DIS3 and AID interaction in CH12F3-2A as well as the signal in the 2 separate background controls performed. The number of interaction sites (PLA spots) in a cell was quantified and demonstrated to be significantly lower in background control which confirms the specificity of interaction. The background control giving the highest signal was chosen for further analysis for a most conservative approach. Analysis was restricted to cells with at least one PLA spot identified (PLA-positive cells). Statistics were performed with the Wilcoxon test comparing given background control with the PLA signal for activated cells. The black horizontal lines represent means. (G-J) Analysis of the distribution of relative distance of regions translocating in murine plasmacytomas to the ChIP-seq peaks active H3K4me1 or H3K4me3 (G) Super enhancers (H) p300 (I) and CTCF (J). Statistics were performed with the Pearson's $\chi^2$ test comparing to observed relative distance frequencies to the expected random distribution.

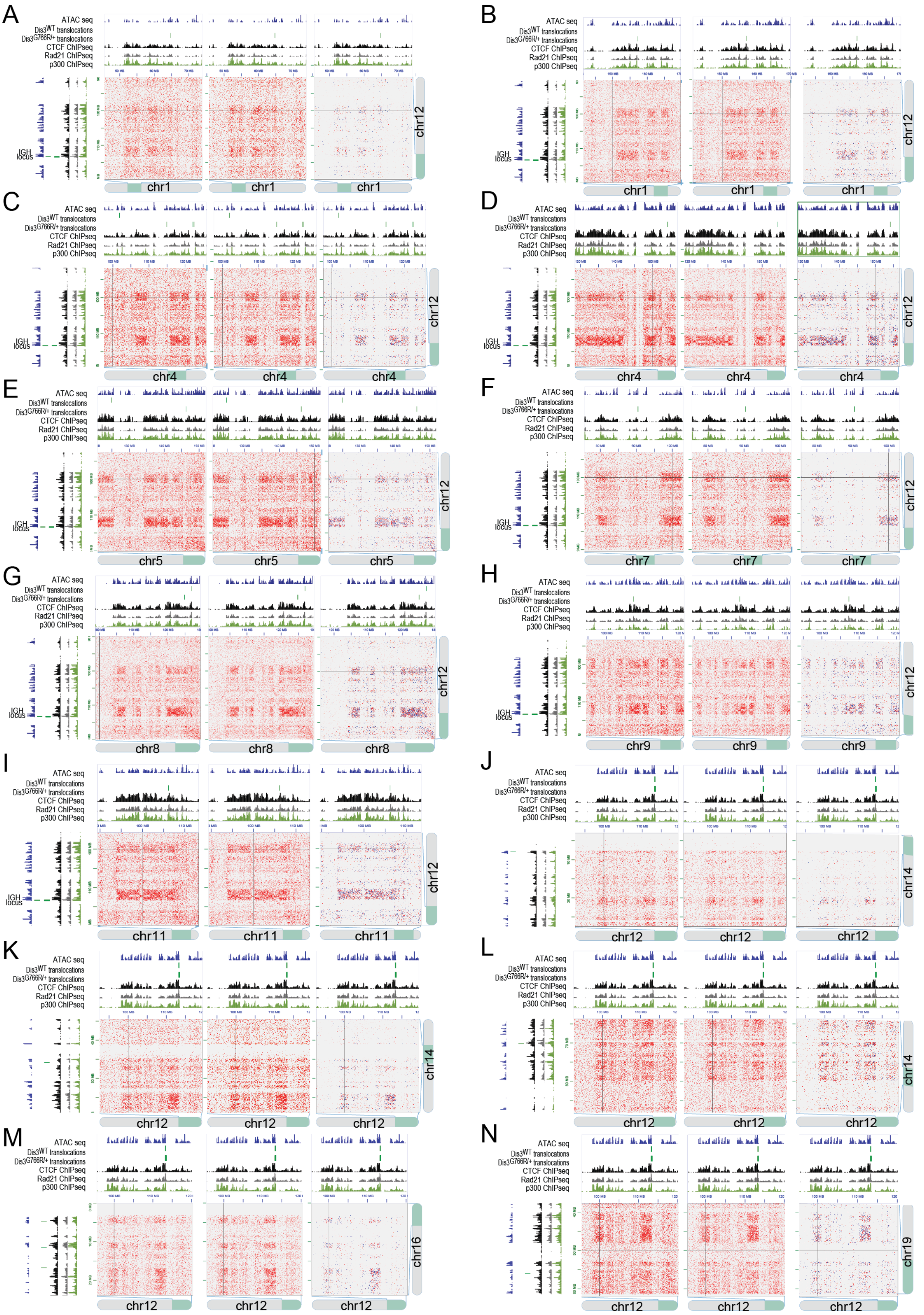


**Figure S7.** The chromatin state of all regions translocating to the IGH locus in murine plasmacytoma of both genotypes. ATAC seq signal and CTCF, Rad21 and p300 occupancy by ChIP seq as well as interchromosomal interactions detected using MicroC of regions translocating from the IGH locus to: chromosome 1(A,B), chromosome 4 (C,D), chromosome 5 (E), chromosome 7 (F), chromosome 8 (G), chromosome 9 (H), chromosome 11 (I), chromosome 14 (J,K,L), chromosome 16 (M) , chromosome 19 (N).

**Description of Supplementary Datasets**

**Table S1.** Exel file. Contains results of differential gene expression analysis from DIS3 G766R vs WT naïve B-cells.

**Table S2.** Exel file. Contains results of differential gene expression analysis from DIS3 G766R vs WT B-cells 3 days post activation.

**Table S3.** Exel file. Contains results of differential gene expression analysis from DIS3 G766R vs WT B-cells 5 days post activation.

**Table S4.** Exel file. Contains results of differential gene expression analysis from DIS3 WT B-cells naïve vs.3 days post activation.

**Table S5.** Excel file. Contains results of differential gene expression analysis from DIS3 G766R B-cells naïve vs. 3 days post activation.

**Table S6.** Excel file. Contains results of differential gene expression analysis from DIS3 WT B-cells naïve vs. 5 days post activation

**Table S7.** Excel file. Contains results of differential gene expression analysis from DIS3 G766R B-cells naïve vs. 5 days post activation.

**Table S8**. Excel file. Contains results of differential protein steady-state levels analysis from DIS3 G766R vs WT B-cells 3 days post activation

**Table S9.** Exel file. Contains results of differential gene expression analysis from DIS3 G766R vs WT MEF cells.

**Table S10.** A bedpe format file with significant chromatin interactions from a MicroC experiment form DIS3$^{G766R/+}$ day3 activated B-cells.

**Table S11.** A bedpe format file with significant chromatin interactions from a MicroC experiment form DIS3 WT day3 activated B-cells.

**Table S12.** A bedpe file format with DIS3$^{G766R/+}$ plasmacytoma translocations.

**Table S13.** A bedpe file format with DIS3 WT plasmacytoma translocations.

**Data, code and materials availability**

The sequencing data discussed in this publication have been deposited in NCBI's Gene Expression Omnibus and are accessible through GEO Series accession number: GSE155631.

The mass spectrometry proteomics data have been deposited to the ProteomeXchange Consortium via the PRIDE repository with the dataset identifier PXD050438.

Any other data, materials and additional information required to reanalyze the data reported in this paper is available from the lead contact upon request
