## Supplementary material for "Multiple Myeloma associated DIS3 mutations drive AID-dependent IGH Translocations": Methods section

**STAR★Methods**

Key resources table

| **REAGENT or RESOURCE** | **SOURCE** | | **IDENTIFIER** |
| --- | --- | --- | --- |
| **Antibodies** | | | |
| Goat anti-Mouse IgG1 | Invitrogen | Cat. No. A-21127, RRID:AB_2535769 | |
| anti-AID | Invitrogen | Cat. No. PA5-18913, RRID:AB_10978944 | |
| anti-DIS3 | Proteintech | Cat. No. 14689-1-AP RRID:AB_2091025 | |
| anti-γH2A.X | Abcam | Cat. No. ab2893, RRID:AB_303388 | |
| Goat anti-Rabbit IgG (H+L) AF555 | Invitrogen | Cat. No. A-21429, RRID:AB_2535850 | |
| anti-CD40 antibody | Invitrogen | Cat. No. MA1-81395, RRID:AB_928280) | |
| **Chemicals, peptides, and recombinant proteins** | | | |
| Il4 | Peprotech | | Cat. No. 214-14 |
| LPS | Santa Cruz Biotechnology | | Cat. No. sc-3535 |
| TGF-β1 | Peprotech | | Cat. No. 100-21 |
| CellTrace™ Yellow Cell Proliferation Kit | Invitrogen | | Cat. No. C34567 |
| LIVE/DEAD™ Fixable Violet Dead Cell Stain Kit | Invitrogen | | Cat. No. L34964 |
| Hoechst 33342 | Invitrogen | | Cat. No. H3570 |
| Propidium Iodide | Sigma-Aldrich | | Cat. No. P4170 |
| Pristane | Sigma-Aldrich | | Cat. No. P2870 |
| DSG | Thermo Scientific | | Cat. No. A35392 |
| Ribonuclease H (RNase H) | NEB | | Cat. No. M0297S |
| **Experimental models: Cell lines** | | | |
| CH12F3 murine B-cell lymphoma cell line | Ricken BRC cell bank | | RRID:CVCL_E068 |
| **Experimental models: Organisms/strains** | | | |
| C57BL/6J | Institut clinique de la souris (ICS) Illkirch, France | |  |
| BALB/CanNCrlCmd | Mossakowski Medical Research Institute, Warsaw, Poland | |  |
| C57BL/6J-BS-K706-tm1c | Institut clinique de la souris (ICS) Illkirch, France | | IR00003831 |
| C57BL/6J-Dis3em1Iimcb/Tar | This paper | | Dis3em1Iimcb/Tar |
| **Recombinant DNA** | | | |
| pLgw V5-EcoDam | Addgene | | RRID:Addgene_59210 |
| pLgw DIS3_WT-V5-EcoDam | This paper | |  |
| pLgw DIS3_G766R-V5-EcoDam | This paper | |  |
| **Software and algorithms** | | | |
| R | https://www.r-project.org/ | | RRID:SCR_001905 |
| RStudio Workbench | RStudio | | RRID:SCR_000432 |
| BEDTools | https://bedtools.readthedocs.io | | RRID:SCR_006646 |
| SAMtools/BCFtools | https://samtools.github.io | | RRID:SCR_005227 |
| SVDetect | http://svdetect.sourceforge.net/ | | RRID:SCR_010812 |
| STAR | https://github.com/alexdobin/STAR.git | | RRID: SCR_004463 |
| mmsig | https://github.com/evenrus/mmsig.git | |  |
| DESeq2 | 10.18129/B9.bioc.DESeq2 | | RRID:SCR_015687 |
| PON-P2 | http://structure.bmc.lu.se/PON-P2/ | | biotools:pon-p2 |
| MACS2 | https://github.com/macs3-project/MACS | | RRID:SCR_013291 |
| BWtool | https://github.com/CRG-Barcelona/bwtool/wiki | | RRID:SCR_003035 |
| **Other** | | | |
| CoMMpass study high-throughput sequencing data | dbGaP | https://www.ncbi.nlm.nih.gov/gap/ | |

Resource availability

**Lead contact**

Further information and requests for resources and reagents should be directed to and will be fulfilled by the Lead Contact, Andrzej S. Dziembowski.

**Data, code and materials availability**

The sequencing data discussed in this publication have been deposited in NCBI's Gene Expression Omnibus and are accessible through GEO Series accession number: [GSE155631](https://www.ncbi.nlm.nih.gov/geo/query/acc.cgi?acc=GSE155631).

The mass spectrometry proteomics data have been deposited to the ProteomeXchange Consortium via the PRIDE repository with the dataset identifier PXD050438.

Any other data, materials and additional information required to reanalyze the data reported in this paper is available from the lead contact upon request

**Ethical issues**

All procedures were approved by the First Local Ethical Committee in Warsaw affiliated at the University of Warsaw, Faculty of Biology (approval numbers WAW/092/2016, WAW/177/2016, WAW/642/2018). Housing in animal facilities was performed in conformity with local and European Commission regulations under the control of veterinarians and with the assistance of trained technical personnel.

**Method details**

***CH12F3 cell line***

The mouse CH12F3 B-cell lymphoma cell line was acquired from the Ricken BRC cell bank. Cells were cultured in RPMI 1640 ATCC’s modified medium (Invitrogen; A1049101) supplemented with 10% FBS (Invitrogen; 10270-106), 5% NCTC-109 medium (Gibco A/Thermo; 21340039), 50 μM 2-mercaptoethanol, penicillin/streptomycin (Sigma; P4083) at density between 0.5-1 x 10^6^ cells/ml. Media were changed every 3 days.

To obtain lentiviral constructs for the DamID experiment, mouse WT and S766R DIS3 coding sequences were fused to the Dam methylase coding sequence. WT DIS3 cDNA was amplified and cloned into pLgw V5-EcoDam lentiviral constructs (Addgene; 59210) using SLIC. The S766R DIS3 construct was obtained by site-directed mutagenesis.

For the negative control construct, mEGFP was amplified using PCR and cloned into pLgw V5-EcoDam lentiviral constructs (Addgene; 59210) using SLIC. Lentiviral transductions of CH12F3 cells were carried out exactly as described previously. Transduced cells were selected using Zeocin at a concentration of 250 µg/ml for 14 days and subsequently, cell cultures were expanded and preserved in 50% FBS, 40% RPMI medium and 10% DMSO.

CH12F3 cells were activated for 3 days by supplementing the standard culture medium with 1 ng/ml TGF-β1 (Peprotech, 100-21), IL-4 (Peprotech, 214-14)5 ng/ml, 1 μg/ml 𝛂CD40 .

| Primers: |  |
| --- | --- |
| WT DIS3 cDNA F | aagtatatatttttaataagggtgggcgcgatgctcaggtccaagacgttc |
| WT DIS3 cDNA R | cactttgtacaagaaagctgggtcggcgcgcttctcaagcttcctcttc |
| site-directed mutagenesis F | catcactatCgcttagcctccccc |
| site-directed mutagenesis R | gggggaggctaagcGatagtgatg |
| mEGFP F | tccgaaaacctgtacttccaaggaaccggtatggtgagcaagggcgagg |
| mEGFP R | atcaccctgaaaatacaaattctcgctagccttgtacagctcgtccatgc |

***DamID-seq protocol***

The DIS3 DamID-seq procedure and library preparations were performed as described previously ^1^. In short: DNA from CH12F3 cells expressing DIS3 DamID constructs or not, as the control, were isolated using a Bioline Isolate II genomic DNA kit (BIO-52067). First, 0.5 µg of DNA was digested with DpnI (NEB; R0176) in 10 µl volume at 37**°**C for 16h. Next, DpnI was heat-inactivated, and the DpnI adaptor was ligated in 20 µl volume with 2.5 units of T4 DNA ligase (NEB; M0202). This was followed by DpnII digestion in a total volume of 50 µl with 10U of DpnII (NEB; R0543). DNA was amplified by PCR with Taq polymerase using the following program: samples were denatured at 95 for 8 minutes, then amplified for 19 cycles, with: 20’’ of denaturation at 94℃, 30’’ of annealing at 58℃ and 20’’ elongation 72℃ followed by a 2 min incubation at 72℃.

PCR products were cleaned up using a Bioline Isolate II PCR and Gel kit according to the manufacturer’s instructions (BIO-52060). DNA was repaired with Blunt-end ending 3′ overhang using End-It DNA End-Repair kit in 25 µl volume (Epicentre; ER81050), and 3′-A overhang was added with Klenow fragment (NEB; M0212) and dATP. This was followed by purification using CleanPCR beads (CleanNA; CPCR-0005). DNA concentration was measured by Nanodrop and adjusted to 40 ng/µl. Illumina Y-adaptor was ligated, and the libraries were PCR amplified using standard protocols and sequenced with an average read number of 40M.

| Primers: |  |
| --- | --- |
| DpnI PCR | Nnnngtggtcgcggccgaggatc |
| Adaptor_top | ctaatacgactcactatagggcagcgtggtcgcggccgagga |
| DpnI adaptor bottom | Tcctcggccgcg |
| Y-adaptor top | Acactctttccctacacgacgctcttccgatct |
| Y-adaptor bottom | P-gatcggaagagcacacgtct (5′- phosphorylated) |
| P5- Illumina-2 PCR | aatgatacggcgaccaccgagatctacactctttccctacacgacgctcttccgatct |

***PLA procedure***

In situ Proximity Ligation Assay was performed on activated CH12F3-2A cells and non- activated (naive) cells as a control. Activation was performed by seeding cells at 105 cells/ml in culture medium supplemented with 1 μg/ml anti-CD40 antibody (Invitrogen, MA1-81395), 5 ng/ml IL-4 (Peprotech, 214-14) and 1 ng/ml TGF-β1 (Peprotech, 100-21), and cells were grown for 72 h at 37°C in 5% CO2. For each experimental condition, 2×106 cells were spun (400 rcf, 4 min), washed with PBS and spun again. Next, cells were fixed with 3,7% formaldehyde in PBS for 25 min at room temperature, washed with PBS twice, permeabilized with 1% Tween-20 (v/v) in PBS for 15 min at room temperature and washed with PBS once. Then, cells were blocked with 3% BSA in PBS for 1 h at room temperature. In order to reduce Fc receptor-mediated binding by antibodies used in PLA procedure, Mouse BD Fc Block (BD Biosciences, 553142) was added to cells in amount of 1 μg/10^6^ cells and incubated according to manufacturer’s instructions. Primary anti-AID (1:100; Invitrogen, PA5-18913) and anti-DIS3 antibodies (1:100; Proteintech, 14689-1-AP) were added to cells in blocking solution and incubated overnight at 4°C. Additionally, three background controls were performed in which cells were incubated only with anti-AID antibody or only with anti-DIS3 antibody, or without primary antibodies. Next, cells were washed with PBS twice and the in-situ PLA was performed with the Duolink In Situ PLA kits (Sigma-Aldrich) according to manufacturer instructions with the exception that the reactions and washings were carried out in 1,5 ml microcentrifuge tubes, and all the incubations were performed in thermoblock. The interaction between AID and DIS3 was detected using Duolink In Situ PLA Probe Anti-Goat MINUS kit (Sigma-Aldrich, DUO92006) and Duolink In Situ PLA Probe Anti-Rabbit PLUS kit (Sigma-Aldrich, DUO92002) against anti-AID antibody and anti-DIS3 antibody, respectively. Fluorescent signal was generated using Duolink In Situ Detection Reagents Red kit (Sigma-Aldrich, DUO92008). Cells were mounted and the nuclei were stained using the Duolink In Situ Mounting Medium with DAPI (Sigma-Aldrich, DUO82040) as follows: after the last wash, cells were spun, all the supernatant was removed carefully, cells were resuspended thoroughly but gently in 12 µl of Mounting Medium, the cell suspension was dripped onto a microscope slide, covered with a coverslip (Ø 18 mm), and gently pressed down. Edges of the coverslip were sealed with nail polish.

***γH2A.X immunofluorescent staining***

DNA double-strand breaks were stained in mouse B cells using γH2A.X as a marker. For each experimental condition, 10^6^ cells were spun (400 rcf, 4 min), washed with PBS and spun again. All the following steps were carried out in suspension in 1,5 ml microcentrifuge tubes. Washed cells were fixed with 3,7% formaldehyde in PBS for 25 min at room temperature, washed with PBS twice, permeabilized with 1% Tween-20 (v/v) in PBS for 15 min at room temperature and washed with PBS once. Then, cells were blocked with 3% BSA in PBS for 1 h at room temperature. Primary anti-γH2A.X (1:400; Abcam, ab2893) was added to cells in blocking solution and incubated overnight at 4°C. After being washed twice with PBS, cells were incubated with goat anti-rabbit IgG secondary antibody conjugated to Alexa Fluor 555 (1:800; Invitrogen, A-21429) for 1 h at room temperature. Next, cells were washed with PBS once and the nuclei were stained using 2 μg/ml Hoechst 33342 (Invitrogen, H3570) in PBS for 5 min at room temperature. After being washed twice with PBS, cells were spun, all the supernatant was removed carefully, cells were resuspended thoroughly but gently in 12 µL of ProLong Glass Antifade Mountant (P36980, Invitrogen), the cell suspension was dripped onto a microscope slide, covered with a coverslip (Ø 18 mm), and gently pressed down.

***Microscopic analysis***

For Fig. 5, cells were imaged using Zeiss LSM800 confocal microscope with 40×/1.3 oil immersion apochromatic objective and GaAsP PMTs detectors. ZEN software (Zeiss, version 2.6) was used for microscope control. Imaging was performed at a controlled temperature of 22⁰C. Z-stack images were processed using Fiji/ImageJ software (version 1.53f51). For each Z-stack, the maximum intensity Z-projection was applied and then images of PLA channel were subjected to rolling-ball background subtraction with a radius set to 20 pixels and contrast enhancement with 0.025% saturated pixels.

For Fig.5 and Figure EV6, cells were imaged using Olympus IX81-ZDC wide-field fluorescence microscope with 40×/0.95 apochromatic objective and EM-CCD camera (Hamamatsu, C9100-02) equipped with ScanR modular imaging platform (Olympus). ScanR Acquisition software (version 2.2.0.8) was used for microscope control. Imaging was performed at a controlled temperature of 22⁰C. Image analysis was performed using ScanR Analysis software (version 3.0.0). Images were subjected to rolling-ball background subtraction with a radius set to 40 and 2 pixels for DAPI and PLA channel, respectively. Next, segmentation was performed on the DAPI channel using the Intensity algorithm. The identified objects were gated based on a scatter plot of the circularity factor relative to the area. The object that comprised the gate was classified as normal, single nucleus. PLA spots in a cell were searched for as sub-objects in the area encompassing a nucleus and a 7-pixel radius outside it, basing on PLA channel and using the Intensity algorithm. Each identified sub-object was assumed to be a PLA spot. Further numerical data processing and statistical analysis were performed using R environment. Cells containing at least one PLA spot were filtered out and for each experimental condition, a total number of spots in a cell and its mean were calculated. Statistical significance was calculated using the Wilcoxon test. All data were checked beforehand for normality using the Shapiro-Wilk test.

For Fig. 5F,G cells were imaged using Olympus IX81 wide-field fluorescence microscope with 60×/1.35 oil immersion apochromatic objective and CCD camera (Hamamatsu, Orca-R2/C10600) equipped with ScanR modular imaging platform (Olympus). ScanR Acquisition software was used for microscope control. Imaging was performed at a controlled temperature of 22⁰C. Image analysis was performed using ScanR Analysis software (version 3.0.0). Images were subjected to rolling-ball background subtraction with a radius set to 75 and 2 pixels for Hoechst and γH2A.X channel, respectively. Next, segmentation was performed on the Hoechst channel using the Intensity algorithm. The identified objects were gated based on a scatter plot of the circularity factor relative to the area. The object that comprised the gate was classified as normal, single nucleus. γH2A.X foci in a cell were searched for as sub-objects in the area encompassing a nucleus basing on γH2A.X channel and using the Intensity algorithm. Each identified sub-object was assumed to be a DNA double-strand break. Further numerical data processing and statistical analysis were performed using R environment. For each experimental condition a total number of DSB in a cell was calculated. Statistical significance was calculated using the Wilcoxon test.

For Fig. 5F, representative images obtained by the abovementioned imaging procedure were selected. Images were processed using Fiji/ImageJ software (version 1.53f51). Images of both channels were cropped to provide a more accurate visualization of γH2A.X foci in the cells. Images of γH2A.X channel were subjected to rolling-ball background subtraction with a radius set to 3 pixels and contrast enhancement with 0.01% saturated pixels. Images of Hoechst channel were subjected to rolling-ball background subtraction with a radius set to 40 pixels

***Mouse line generation***

The constitutive knock-in G766R mice were constructed according to the strategy described in Figure EV4A in the Institut clinique de la Souris (ICS) Illkirch, France using homologoues recombination technology in mES cells in the C57BL/6N background.

In brief, embryonic stem (ES) cells were cultured following the standard protocol ^2^ and then electroporated with the targeting vector. This vector carried a G766R point mutation introduced into exon 17, specifically a G to C substitution at position 2393 of the gene sequence. The mutation site was flanked by 4 and 3.5 kb homology arms (Figure EV1B) and accompanied by a floxed CMV promoter-driven Neo-cassette expressing Cre-ER, which was inserted into the intron of the Dis3 gene. Following targeting, ES cells were selected using G418, and clones were genotyped to identify those carrying the Dis3G766R allele. These cells were treated with tamoxifen to remove the cassette, leaving a single loxP site in intron 19. ES cells carrying the Dis3G766R allele were subsequently injected into C57BL/6N blastocysts, which were then implanted into the uterine horns of pseudopregnant C57BL/6N mice. Chimeras and their subsequent progeny were mated with C57BL/6N wild-type mice to generate germline production and expand the colony. The progeny resulting from this cross-breeding produced heterozygous mice Dis3G766R/+ as well as wild-type (WT) controls.

Mice are genotyped by PCR using primers:

| 766seq_F | gtctcaaaaacacacggcagagactc |
| --- | --- |
| 766seq_R | gaggatgtgtgagataagatacaccgg |

Dis3 WT allele migrates as a 202 bp band, whereas the mutant G766R allele gives a 281 bp amplicon.

***Mice breeding conditions***

Mice were bred in the animal house of Faculty of Biology, University of Warsaw maintained under conventional conditions in a room with controlled temperature (22 ± 2°C) and humidity (55 ± 10%) under a 12 h light/12 h dark cycle in open polypropylene cages filled with wood chip bedding (Rettenmaier). The environment was enriched with nest material and paper tubes. Mice were fed ad libitum with a standard laboratory diet (Labofeed B, Morawski). Animals were closely followed-up by the animal caretakers and researchers, with regular inspection by a veterinarian, according to the standard health and animal welfare procedures of the local animal facility.

***Mouse embryonic fibroblast primary cell cultures***

Primary cell cultures of mouse embryonic fibroblasts (MEF) were isolated from E13.5 embryos as defined by the date of the vaginal plug and confirmed by characteristic features of the embryo morphology. Embryo heads were used for DNA isolation and genotyping. The corpses were gutted and mechanically disintegrated by chopping and passing the tissue fragments suspended in culture media (DMEM supplemented with 10% FBS, 50 μM 2-mercaptoethanol, penicillin/streptomycin) with a syringe through the gauge 19 needles. Disintegrated tissue fragments of a single individual were left to settle in 100 mm petri dishes and form outgrowths. After 3 days, cells were treated with 0.25% Trypsin in 1 mM EDTA for 3 min. Cells were passaged for at least 3 passages before they were collected or used for further applications. Cells were passaged every 3 days. Cells isolated from 3 independent litters in pairs, including DIS3^+/+^ and DIS3^G766R/+^ were used for the RNA-seq library preparation.

For the 3T3 protocol, MEF cells from 12 DIS3^+/+^ and 13 DIS3^G766R/+^ littermates were used. Cells passaged 2 times to disintegrate into a single cell suspension and noted. Cells were plated into a 10-cm cell culture dish at a density of 9 × 10^5^ cells per dish. Cells were cultured at standard conditions with medium change after 2 days in culture. Three days after plating, cells were trypsinized, counted using a hemocytometer, and replated 9 × 10^5^ into a 10-cm cell culture dish. The strict passaging regime was repeated for 30 passages or until the immortalization of cells was obtained. Cumulative cell numbers were plotted.

***Pristane-induced mouse plasmacytoma (MPC) model***

Six- to ten-week old mice were injected 3 times intraperitoneally, in 2-month intervals, with 0.5 ml of pristane oil (2,6,10,12-tetramethylpentadecane) (Sigma-Aldrich, Cat. No. P2870) to induce the development of mouse plasmacytoma (MPC), an established murine model of multiple myeloma ^3,4^. The day of the first injection is considered as day 0 of the experiment. Mice were regularly observed for abdominal swelling. The neoplastic nature of the observed ascites was confirmed by paracentesis and ascitic smear stained with Wright-Giemsa stain which had to confirm >10 atipical plasma cells. For a limited number of samples the smers were . Animals were closely followed-up by the animal caretakers and researchers, with with particular attention to lupus-like syndrome signes such as alveolar hemorrhage or fur loss. Regular inspection by a veterinarian was conducted, according to the standard health and animal welfare procedures of the local animal facility.

Mice for the pristane-induced mouse plasmacytoma experiment were maintained in individually ventilated cages in a room with controlled temperature (22 ± 2°C) and humidity under a 12 h light/12 h dark cycle.

As plasmacytomas in a C57Bl/6 background are a rare event in the WT genotype, to obtain a sufficient amount of material for and better compare plasmacytomas from Dis3^G766R/+^ and WT mice, we have backcrossed Dis3^G766R/+^ mouse line into plasmacytoma prevalent BALB/C background (more than 6 crosses) and used WT and Dis3^G766R/+^ littermates to induce plasmacytomaTo obtain G766R mice on a BALB/C genetic background, DIS3^G766R/+^ mice were backcrossed to BALB/CanNCrlCmd mice for 6 generations.

***Tissue collection and primary cell culture, in vitro B-cell activation, CSR efficiency assay***

Splenic naïve B-cells were prepared using the CD43 negative selection commercial kit (Miltenyi Biotec.; 130-090-862). Spleens were pulled from 2-3 sacrificed sex- and age-matched animals mechanically disintegrated and filtered through a 70 µm cell strainer to prepare a single cell suspension essential for microbead separation. Spleen extracts were depleted from red blood cells using ACK lysis buffer. The magnetic bead, column-based enrichment of naïve B-cells was performed according to the manufacturer’s instructions. Cells were resuspended and cultured in RPMI 1640 containing 15% FBS supplemented with 100 nM 2-mercaptoethanol and penicillin/streptomycin (Sigma; P4083).

Ex vivo activation of B lymphocytes for class switch recombination, proliferation studies and RNA sequencing and was achieved by culturing them in RPMI 1640 containing 15% FBS supplemented with 100 nM 2-mercaptoethanol, 20 µg/ml LPS and 20 ng/ml IL-4. CSR efficiency was assessed in B-cells 3 days after activation. In brief, cells were collected and stained for 30 min with Alexa Fluor 555 conjugated goat antibody raised against mouse IgG1 (Invitrogen; A-21127) at 10 µg/ml. The percentage of positive cells that underwent IgG1 class switch recombination was quantified using flow cytometry on an Attune NxT Cytometer (Thermo) or CytoFLEX Flow Cytometer (Beckman Coulter) and analyzed using FlowJo (v10.7.1) software.

***CellTrace Proliferation Assay***

CellTrace™ Yellow Proliferation kit was used according to the manufacture’s instructions. Briefly: Cells were stained by adding 1 μl of CellTrace™ Yellow (Invitrogen, Cat #C34567) stock solution in DMSO per each ml of 10^6^/ml cell suspension in PBS (5 μM working concentration), and subsequently incubated for 40 minutes at room temperature, while being protected from light. Afterward, five times the original staining volume of culture medium, containing FBS, was added to the cells and incubated for 5 minutes. The cells were then pelleted and resuspended in a fresh pre-warmed complete culture medium. 10 minute incubation period was allowed before analysis to enable the acetate hydrolysis of the CellTrace™ reagent. Cells were pipetted in technical triplicates, 3x10^5^ per well, into round bottom 96 well plate in 200 μl growth medium. Cells were activated by supplementing media with 20 µg/ml LPS and 20 ng/ml IL-4 and cultured for 3 days. Naïve B-cells and unstained control cells were parallelly cultured in triplicates. Before cytometric analysis, dead cells were stained with LIVE/DEAD fixable dyes (Invitrogen™) according to manufactures instructions and analized on a CytoFLEX Flow Cytometer (Beckman Coulter). Data were analyzed using FlowJo (v10.7.1) software using the Proliferation functionality.

***RNA-seq, Mate-Pair and Paired-End DNA-seq***

Total RNA from collected tissues and primary cultures was isolated using a standard TRIzol protocol followed by ribodepletion with a Ribo-Zero Gold rRNA Removal Kit (Human/Mouse/Rat) according to the manufacturer’s instructions (Epicentre; RZG1224). Strand-specific libraries were prepared using a TruSeq RNA Sample Preparation Kit (Illumina) and the dUTP method. Three independent replicate sample sets were prepared for each condition. DNA was purified using a SherlockAX kit optimized for small samples (A&A Biotechnology; 095-25). NEXTERA XT mate-pair and TruSeq DNA libraries were prepared according to manufacturer's instructions (Illumina; FC-131-1024 and FC-121-2001). All of library types were sequenced using Illumina NextSeq or NovaSeq platform in a paired-end mode with different average sequencing depth: MEF RNA-seq: 29M reads, Mate-pair DNA libraries: 42.7M reads, standard DNA-seq libraries: 281.5M reads (NovaSeq). The total RNA extracted from splenic primary B-cell cultures, naïve and in vitro activated for 72h and 96h has been commercially sequenced by Azenta Life Sciences. They utilized their strand-specific library protocol, achieving an average depth of 23 million 100 nucleotide paired-end reads.

***DRIPcseq (DNA/RNA hybrid immunoprecipitation followed by cDNA conversion and sequencing)***

After 72h h, the culture medium was removed, and cells were washed once with cold PBS and twice with TE buffer (10 mM Tris [pH 8.0] and 1 mM EDTA). Cell lysis was performed directly on the plate by adding 0.5 ml of lysis buffer (TE supplemented with 0.625% SDS and 0.0625 mg/ml 7 Proteinase K), and the suspension was immediately transferred to a fresh 2 ml tube and incubated for 4.5 h at 37°C with gentle mixing every 15 min. Afterward, genomic DNA was purified by phenol:chloroform:isoamyl alcohol (25:24:1) extraction and ethanol precipitation. Precipitated DNA was washed with 70% ethanol, air-dried, and resuspended in 30 µl of TE buffer. Next, pure gDNA was sheared by sonication using Bioruptor Plus with a water-cooling system (Diagenode) as the following: 3 µg gDNA diluted to 200 µl in TE was placed in 1.5 ml DNA LoBind Tubes (Eppendorf) and subjected to 30 cycles of sonication (30 s ON, 90 s OFF) in “Low” mode. The length of fragments after sonication was evaluated by running DNA samples before and after shearing on a 1.5% agarose gel in 1X TAE (40 mM Tris [pH 8.0], 20 mM acetic acid, and 2 mM EDTA) at 70-80 V. To ensure specificity, nucleic acids are pre-treated with Ribonuclease H (RNase H) (NEB, cat. No. M0297S) to eliminate RNA:DNA hybrids from the mixture. In this process, 4 μg of digested gDNA is incubated with 4 μL of NEB RNase H for 4 hours at 37°C. Following this treatment, proceed with the S9.6 immunoprecipitation step, processing both the control, RNase H-treated samples alongside the experimental samples. Afterward, 4 µg of fragmented gDNA was mixed with 10 µg of S9.6 antibody, diluted to a final volume of 1 ml with binding buffer (20 mM HEPES [pH 7.4], 150 mM NaCl, and 0.05% Triton X-100), and incubated overnight with gentle rotation in a refrigeration cabinet. The next day, input fractions that contained S9.6 were added to 50 µl of M2 anti-FLAG Magnetic Beads (Sigma) or Protein G Dynabeads (Thermo), respectively, and incubated for 2 h. Additionally, a second set of negative controls were prepared by incubating 4 µg of sonicated gDNA with beads without the bait. Following incubation, the unbound fraction was removed, and beads were washed three times with 500 µl of binding buffer for 5 min with gentle rotation in a refrigeration cabinet. Elution was performed by adding 125 µl of elution buffer (50 mM Tris [pH 8], 1 mM EDTA, 0.5% SDS, and 0.64 mg/ml Proteinase K) for 45 min at 55°C. Two replicates of each sample were pooled and purified using a DNA clean and concentrator kit (Zymo Research) according to the manufacturer’s protocol. After elution with 30 µl of elution buffer (Zymo Research), half of the eluate was treated with 2 µl of DNAse I (1 U/µl) in 1X DNase I digestion buffer (Thermo Fisher Scientific) for 1 h at 37°C. For DNase I inactivation, 1 µl of 0.5 M EDTA was added for 10 min at 65°C. RNA was purified using the RNA clean and concentrator kit (Zymo Research). The purified RNA underwent NGS library preparation and subsequent sequencing, which was conducted commercially by Azenta Life Sciences using their strand-specific protocol*.*

***MicroC***

MicroC was performed according to ^5^ with minor modifications. Two replicates of B cells, isolated from WT and Het mice, were activated for 72hours by incubation with LPS and IL4 (20 µg/ml LPS and 20 ng/ml IL-4 final concentration) prior to experiment. Next, cells were collected by trypsinization, washed in PBS and counted. For crosslinking, 300 mM DSG (ThermoScientific) solution in DMSO (100x) was prepared freshly and diluted to 1x in room temperature PBS. Cellular pellet was resuspended in DSG solution and incubated for 35 minutes on rotating wheel at room temperature. Next 16% methanol-free formaldehyde (Thermo Scientific) was added to a mixture to a final concentration of 1% and incubated for next 10 minutes on rotating wheel at room temperature. To quench the crosslinking reaction, Tris pH 7.5 was added to a final concentration of 0.375 M and incubated on rotating wheel for 5 minutes at room temperature. Next, cells were centrifuged at 4°C, washed once with ice-cold PBS and re-counted. Cells were divided into 5 mln cells portions, pelleted, snap frozen and stored at -80°C until needed.

Before MicroC protocol, MNase titration was performed to specify the required amount of enzyme for sufficient chromatin cleavage. Next, 10 million crosslinked activated B cells from WT and Het mice were resuspended in MB#1 buffer (10 mM Tris pH 7.5, 50 mM NaCl, 5 mM NgCl2, 1mM CaCl2, 0.1% Igepal, EDTA-free protease inhibitor cocktail (Roche)) and incubated on ice for 20 minutes to isolate nuclei. After one wash with MB#1 buffer, nuclei were resuspended in MB#1 buffer containing MNAse and incubated for 20 minutes in 37°C with 1000 rpm rotation in thermoblock to digest chromatin. Reaction was stopped by addition of 4mM EGTA and 10 minutes incubation at 65°C. Next, nuclear pellet was washed twice in MB#2 buffer (10 mM Tris pH 7.5, 50 mM NaCl, 10 mM MgCl2, BSA), resuspended in 100 ul Repair End Master Mix (1x NEB 2.1, 2mM ATP, 5mM DTT, 50 U PNK), incubated at 37°C for 15 mins with shaking 1000 rpm. Next, 50 U Klenow was added and incubated for next 15 mins at 37°C with shaking 1000 rpm. To biotinylate, Biotin Master Mix (66 nM Biotin-dATP, 66 nM Biotin-dCTP, 66 nM dTTP, 66 nM dGTP, T4 DNA ligase buffer, 33 ug BSA) was added and incubated 45 mins 25°C with shaking 1000 rpm. To quench the reaction EDTA was added to final concentration of 30 mM, incubated at 65°C for 20 minutes and the chromatin pellet was washed once with MB#3 buffer (50 mM Tris pH 7.5, 10 mM MgCl2, 50 μg BSA). Next, the chromatin pellet was resuspended in Ligation Master Mix (T4 buffer, 100 μg BSA, 10 000 U T4 DNA ligase), incubated for 2.5 hours at 25°C and centrifuged. The chromatin pellet was resuspended in Exonuclease Master Mix (NEBuffer 1, 1000 U Exonuclease III), incubated for 15 mins at 37°C and centrifuged. Next, the samples were incubated overnight at 65°C in Reverse crosslinking solution (1% SDS, 200 mM NaCl, 520 ug Proteinase K, 26 ug RNAseA), DNA was purified on 1.0x AmPure beads to enrich for dinucleosome fraction, concentration was measured and aliquot was checked on agarose gel to ensure efficient MNase cleavage.

Prior to library preparation 10-15ug biotinylated chromatin was pull-down using Dynabeads MyOne Streptavidin T1 for 30 mins incubation at room temperature, washed twice with 1xTWB (5 mM Tris-HCl pH 7.5, 0.5 mM EDTA pH 8.0, 0.1% Tween-20) and once with 10 mM Tris pH 7.5.

Library was prepared using NEBNext® Ultra™ II DNA Library Prep Kit for Illumina® according to manufacturer protocol. Briefly, beads were resuspended in 25 ul water, 3.5 ul End Prep Reaction Buffer and 1.5 ul End Prep Enzyme Mix and incubated in thermocycler for 30 mins at 20°C, followed by 30 mins at 65°C. Next, 15 ul Ligation Master Mix, 0.5 ul Ligation Enhancer and 2.5 ul unique xGen UDI-UMI Adapter with indexes (IDT) were added and incubated for 30 mins at 20°C. The beads were washed twice with 1xTWB and once with 10 mM Tris pH 7.5. Test PCR using Q5 NEB Mix was run on 10% beads to establish required number of PCR cycles and final PCR was performed accordingly. Eventually, PCR product was purified with two-sided AmPure size selection (0.6x and 1x), measured using Qubit and checked on TapeStation. The libraries were paired-end 150 bp sequenced to around 300 MR each.

***Global proteomics by liquid chromatography (LC)-MS***

Cell pellets were subjected to the Sample Preparation by Easy Extraction and Digestion (SPEED) protocol^6^. In brief, cells were solubilized in concentrated TFA and incubated for 10 minutes at RT. Samples were neutralized by adding 2 M Tris-Base buffer using 10×volume of TFA and further incubated at 95°C for 5 min after adding Tris(2-carboxyethyl)phosphine (TCEP) to a final concentration of 10 mM and 2-Chloroacetamide (CAA) to a final concentration of 40 mM. Protein concentrations were determined by turbidity measurements at 360 nm, adjusted to the same concentration using a sample dilution buffer (2M TrisBase/TFA 10:1 (v/v)) and then diluted 1:4-1:5 with water. Digestion was carried out overnight at 37°C using trypsin at a protein/enzyme ratio of 100:1. TFA was added to a final concentration of 2% to stop digestion. The resulting peptides were TMT labelled on-StageTip. TMT-labelled samples were compiled into a single TMT6 sample and concentrated using a SpeedVac concentrator. Peptides in the compiled sample were fractionated (6 fractions) using the bRP fractionation. Prior to LC-MS measurement, the peptide fractions were resuspended in 0.1% TFA, 2% acetonitrile in water. Prior to the liquid chromatography (LC)-MS measurement, the peptide fractions were resuspended in 0.1% TFA and 2% acetonitrile in water. Chromatographic separation was performed on an Easy-Spray Acclaim PepMap column (50cm length × 75 µm inner diameter; Thermo Fisher Scientific) at 55°C by applying 90 min acetonitrile gradients in 0.1% aqueous formic acid at a flow rate of 300 nl/min. An UltiMate 3000 nano-LC system was coupled to a Q Exactive HF-X mass spectrometer via an easy-spray source (all Thermo Fisher Scientific). The Q Exactive HF-X was operated in TMT mode with survey scans acquired at a resolution of 60,000 at m/z 200. Up to 18 of the most abundant isotope patterns with charges 2-5 from the survey scan were selected with an isolation window of 0.7 m/z and fragmented by higher-energy collision dissociation with normalized collision energies of 32, while the dynamic exclusion was set to 35 s. The maximum ion injection times for the survey scan and dual MS (MS/MS) scans (acquired with a resolution of 45,000 at m/z 200) were 50 and 130 ms, respectively. The ion target value for MS was set to 3e6 and for MS/MS was set to 1e5, and the minimum AGC target was set to 1e3.The data were processed with MaxQuant v. 1.6.17.0 ^7^ and the peptides were identified from the MS/MS spectra searched against Uniprot mouse reference proteome (UP000000589) using the built-in Andromeda search engine. Reporter ion MS2-based quantification was applied with reporter mass tolerance = 0.003 Da and min. reporter PIF = 0.75. Cysteine carbamidomethylation was set as a fixed modification and methionine oxidation, glutamine/asparagine deamination, as well as protein N-terminal acetylation, were set as variable modifications. For in silico digests of the reference proteome, cleavages of arginine or lysine followed by any amino acid were allowed (trypsin/P), and up to two missed cleavages were allowed. The false discovery rate (FDR) was set to 0.01 for peptides, proteins, and sites. A match between runs was enabled. Other parameters were used as pre-set in the software. Unique and razor peptides were used for quantification enabling protein grouping (razor peptides are the peptides uniquely assigned to protein groups and not to individual proteins). Reporter intensity corrected values for protein groups were loaded into Perseus v. 1.6.10.0^8^. Standard filtering steps were applied to clean up each dataset: reverse (matched to decoy database), only identified by site, and potential contaminant (from a list of commonly occurring contaminants included in MaxQuant) protein groups were removed. Reporter intensity corrected values were Log2 transformed. Protein groups with all values were kept. Reporter intensity values were then normalized by median subtraction within TMT channels. Student’s t-tests were performed on the these values for groups of samples constituting a given dataset. Student’s t-test (permutation-based FDR = 0.01, S0 = 0.1) was performed to return protein groups, which levels were statistically significantly changed between the sample groups investigated. This dataset has been deposited to the ProteomeXchange Consortium via the PRIDE repository with the dataset identifier PXD050438.

***Bioinformatic analysis***

MMRF CoMMpass study whole-genome DNA sequencing reads were mapped to the GRCh38 human reference genome, using STAR^9^ processed and filtered using SAMtools^10^. To identify translocations, significant structural variants were called using SVDetect^11^ with default settings. Significant translocations were filtered for those known to be commonly initiated in the Igh locus and genotype. RNA sequencing reads were mapped using STAR split read aligner. RNA sequencing samples of patients genotyped based on exome sequencing as being a DIS3 somatic mutation or a recurrent DIS3 mutation were compared to RNA samples from patients not possessing such a genotype as well as not possessing a germline DIS3 mutation or a mutant DIS3 variant as a minor clone. Reads were counted onto a gene annotation (gencode v34 basic) using FeatureCount (subread package release 2.0.0)^12^. Differential expression RNA-seq analyses were performed as previously using DESeq2 R package^13^. Whole-exome DNA sequencing reads were mapped to the GRCh38 human reference genome, using STAR split read aligner^9^, and quantified using SAMtools and BCFtools pipeline^10^. Healthy tissue control whole-exome DNA sequencing data were mapped and variants that were identified in at least one control sample were defined as germline mutations. The Single Base Substitution (SBS) profiles were calculated using SigProfilerMatrixGenerator^14^ and fitted to the COSMIC Mutational Signatures (v3.2) using mmsig^15^.

To identify translocations in plasmacytoma DNA samples, mate-pair DNA sequencing data were preprocessed with NXtrim^16^. Both mate-pair and standard TruSeq DNA libraries were mapped by STAR split read aligner. Statistically significant structural variants were called using SVDetect with default settings. All significant structural variants from all sequencing experiments have been concatenated together according to the genotype and merged using bedtools merge to exclude loci involved in more than one structural variant to be analyzed multiple times. These annotations have been used for the meta-analysis. For the filtering to identify those involving an extended region, including the immunoglobulin heavy chain locus, custom scripts were used.

DamID-seq data were stripped of sequencing adapters using cutadapt and mapped to the mouse reference genome (GRCm38) using STAR aligner. Uniquely mapping reads were counted onto DpnI restriction fragments in the mm10 genome. Count tables were library size corrected. Counts for the 3 replicates of the DamOnly negative control have been averaged and subtracted from all replicates to eliminate the background signal. The annotation of putative PROMPT 3kb regions upstream and antisense to the known transcription start sites in the gencode release M25. Mean values have been calculated of DIS3 occupancy.

ChIP-seq reads were aligned to the mouse reference genome (GRCm38.p6) using bowtie2. All replicates were pooled and used to call peaks with MACS2 (v2.2.7.1) using IGG control as the background. The obtained tracks and peak annotations were utilized in downstream analysis. Super-enhancer annotations were obtained from publicly available data (GSE6206326)^17^. The coordinates, originally in the mm9 genome assembly, were converted to the GRCm38.p6 genome assembly using the UCSC Genome Browser's liftOver tool.

Meta-analyses of genomic regions were performed using custom scripts employing BWtool. Structural variants hotspots were tested for nucleotide composition similarities and their relation to AID chromatin occupancy or transcription using custom scripts. The AID DNA recognition motifs tracks were prepared by calculating the density of RGYW, GAGCT, GGGGW, and GGGCT sequence occurrence in the GRCm38.p6 mouse reference genome. Two G-quadruplex prediction annotations were performed by scanning the GRCm38.p6 mouse genome for at least 4 stretches of at least three G separated by up to 7 of any nucleotides for a more stringent annotation or separated by 30 of any nucleotides for the less stringent annotation. From these annotations, strand-specific density tracks were prepared and used for meta-analysis.

To assess the spatial relationship between chromosomal translocation breakpoints and genomic features associated with chromatin activity, we compared their distances to significant ChIP-seq peaks (H3K4me1, H3K4me3, CTCF, Rad21, p300) and super-enhancer annotation. The analysis was conducted using BEDTools. For each breakpoint, the nearest chromatin feature was identified using bedtools closest. Randomized control datasets were generated by shuffling breakpoint coordinates across the genome while preserving chromosomal distribution, using bedtools shuffle. The absolute distances between breakpoints and features in the observed and randomized datasets were compared using the Wilcoxon rank-sum test to assess enrichment or depletion of proximity.

Additionally, the relative distance distributions (fractional distances within feature intervals) were calculated using bedtools reldist. Observed relative distance frequencies were compared to those expected under the random model using Pearson's χ² test.

The likelihood of G766R mutation pathogenicity, was assessed using the PON-P2 algorithm^18^, with default settings. This machine learning-based classifier categorizes variants into pathogenic, neutral, and unknown classes by utilizing various information, such as evolutionary conservation of sequences, physical and biochemical properties of amino acids, GO annotations, and, if accessible, functional annotations of variation sites.

Raw Hi-C reads were trimmed using TrimGalore (v0.6.7)^19,20^. The fastq files were processed using the Juicer Pipeline (v2.13.07)^21^, using default options and aligned to the GRCm38/mm10 genome assembly. The processing was performed without any input for restriction enzyme sites as MicroC uses the Micrococcal Nuclease (MNase) which does not cleave at specific cut-sites.

Raw ligation frequencies were obtained from the .hic files using juicer tool^3^ with the following command: juicer dump observed NONE <input.hic> <chr12> <chr12> BP 2000 output.txt for each replicate separately. Next, the data was imported to R. The ligation frequency matrix for chromosome 12 was normalized using the Iterative Proportional Fit algorithm (matrix balancing) as described previously^22^ (<https://pekowskalab.nencki.edu.pl/research/the-code>).

Publicly available data from the following publications were used: ChIP-seq AID SRA: SRP003605^23^; ChIP-seq H3K4me3, H3K4me1, H3K27ac, p300, CTCF and Rad21 SRA: SRP075985^24^; EXOSC3 KO RNA-seq SRA: SRP042355^25^; GRO-seq B-cells and CH12F3 SRA: SRP193758^26^; Super enhacer annotation: GSE62063^17^.

19.	Krueger, F. TrimGalore! https://www.bioinformatics.babraham.ac.uk/projects/trim_galore/. https://doi.org/10.5281/zenodo.7598955.
